# Dopamine signatures of excessive and compulsive cocaine and fentanyl use

**DOI:** 10.64898/2026.01.18.700215

**Authors:** Ke Chen, Hao Zheng, Gabrielle Sevrain, Wenxi Xiao, Charlotte Wang, Akili Sundai, Ila Fiete, Fan Wang

**Affiliations:** Department of Brain and Cognitive Sciences, Massachusetts Institute of Technology; Cambridge, 02139, USA; McGovern Institute for Brain Research, Massachusetts Institute of Technology; Cambridge, 02139, USA; Yang Tan Collective, Massachusetts Institute of Technology; Cambridge, 02139, USA

## Abstract

Excessive and compulsive drug use despite adverse consequences is a hallmark of substance use disorder, yet individuals differ markedly in their vulnerability to develop these behaviors. Drugs of abuse are long known to alter endogenous dopamine (DA) signaling, but shared principles for how DA dynamics impact compulsive use among individuals and across drug classes are lacking. Here, we monitored DA release in the medial shell of nucleus accumbens (NAc) during cocaine and fentanyl self-administration, with or without coincident punishment, in large cohorts of mice. Contingent cocaine and fentanyl self-administration evoked complex and individually distinct DA dynamics; nevertheless, a robust negative correlation held across both drugs, such that high takers exhibited lower drug-evoked DA signals. During punished drug taking, cocaine and fentanyl cases were associated with distinct DA signatures of compulsivity. For cocaine, punishment-resistant mice showed lower sustained DA responses during the post-shock, drug-associated cue period, whereas for fentanyl, punishment-resistant mice displayed larger phasic DA at the co-occurrence of footshock and drug infusion. To identify common principles underlying these observations, we developed a computational model grounded in an Actor-Critic temporal-difference (TD) learning framework that incorporates internal states, agent’s uncertainty, and drug-specific effects. Remarkably, this model captures the observed diversity in DA dynamics across drug classes and among mice with variable drug taking propensities, hereby providing a unified interpretation of NAc DA signals as encoding TD reward prediction errors.

## Main Text

Substance use disorder (SUD) is a leading cause of drug overdose-related mortality. Excessive drug consumption and compulsive drug taking despite adverse consequences are defining features of SUD. However, only a subset of individuals transition from initially controlled, recreational drug use to uncontrolled and compulsive taking, underscoring a striking degree of individual variability in vulnerability to addiction(*1*–*6*). Rodent models similarly reveal significant individual differences in the propensity to transition from controlled to addiction-like drug use(*7*– *15*). For example, in cocaine self-administration paradigms, only a subset of rats met operational criteria for addiction-like behaviors, characterized by escalation of intake, persistent drug seeking despite adverse consequences, and heightened motivation to obtain the drug(*13*, *15*). Despite extensive research, the neurobiological mechanisms by which exposure to addictive drugs leads to divergent outcomes across individuals remain poorly understood. This study aims to identify specific neurochemical signatures associated with excessive and compulsive drug taking despite adverse consequences.

A large body of work has identified the mesolimbic dopamine system—particularly projections from the ventral tegmental area (VTA) to the nucleus accumbens (NAc)—as central to the reinforcing effect of abused drugs and the development of addiction(*16*–*24*). Seminal microdialysis and voltammetry studies showed that pharmacologically diverse drugs of abuse, including cocaine and opioids, preferentially elevate extracellular dopamine levels in the NAc relative to dorsal striatum in animals(*18*, *23*, *25*). However, dopamine responses to drugs and drug-associated cues are not uniform across individuals, nor do they remain static over the course of addiction development. Indeed, human neuroimaging and rodent studies have revealed substantial inter-individual variability in both drug-evoked and cue-evoked dopamine release(*26*– *30*), suggesting that differences in dopamine responses may contribute to addiction vulnerability(*5*, *16*, *27*, *29*, *31*). From a computational perspective, dopamine has often been conceptualized as a reward prediction error signal within temporal-difference reinforcement learning models(*32*–*35*), whereas psychological and neurobiological theories such as incentive sensitization, allostasis, and habit formation emphasize how drug-induced adaptations in dopamine circuits may drive pathological “wanting”, negative reinforcement, and stimulus–response habits in the development of addiction(*5*, *36*–*40*). Despite these influential theories, direct comparisons of in vivo dopamine dynamics across individuals and across different drug classes remain limited, hindering efforts to determine which dopamine theories best explain experimental data and individual differences in drug taking and compulsive behavior.

In vivo measurements of dopamine using fast-scan cyclic voltammetry (FSCV) and genetically encoded dopamine sensors have provided rich descriptions of dopamine dynamics during psychostimulant self-administration, particularly for cocaine(*23*, *30*, *41*–*43*). In contrast, the subsecond dopamine dynamics underlying opioid self-administration remain poorly characterized, especially in the context of fentanyl. A recent study showed that heroin selectively activates a subset of VTA dopamine neurons projecting to the medial shell of NAc, and that these dopamine neurons are critical for heroin self-administration(*19*). However, little is known about how variability in real-time dopamine dynamics in NAc relates to individual differences in excessive opioid self-administration. Moreover, it remains unknown how dopamine release patterns correlate with the emergence of compulsive drug-taking despite adverse consequences. Critically, few studies have directly compared dopamine signatures across drug classes using the same self-administration paradigm, or tested which formal theories of dopamine in addiction best account for the observed dopamine dynamics across conditions.

Here, we measure real-time changes in dopamine release with a genetically encoded sensor in the medial shell of the NAc as mice develop excessive and compulsive self-administration of two distinct drug types, cocaine and fentanyl. We chose fentanyl to represent the opioid drug class for it being the leading cause of overdose death(*44*), and cocaine to represent the stimulant drug class for its strong abuse potential and for ensuring consistency of our results with existing literature. We focused on the medial NAc shell because this region has been strongly implicated in the reinforcing and motivational effects of both cocaine and opioids(*19*, *41*, *45*, *46*). To capture the transition from controlled to compulsive drug use, we employed long-access intravenous self-administration paradigms, which reliably promote escalation of intake and the emergence of punishment-resistant, compulsive drug taking(*11*, *13*, *47*, *48*). We quantify individual differences in cocaine- and fentanyl-taking behaviors and punishment sensitivity and correlate these measurements with individual variations in dopamine dynamics. We then build a computational model that remarkably recapitulates the diverse dopamine patterns observed across individuals, drug classes, and stages of drug self-administration. Altogether, our study reveals dopamine signatures of excessive and compulsive drug taking in SUD models and provides a unified computational framework for understanding NAc dopamine signals in reinforced drug consumption.

### Individual differences in drug-taking behavior and punishment resistance

To assess individual differences in addiction-like behaviors induced by psychostimulant or opioid exposure, we trained large cohorts of mice to perform intravenous self-administration (IVSA) of either cocaine (n = 24) or fentanyl (n = 27). We also expressed a genetically encoded dopamine (DA) sensor in these mice to measure DA release (described below). Specifically, mice implanted with catheters were trained to press an active lever that triggered either cocaine (0.3 mg/kg/infusion) or fentanyl (2 µg/kg/infusion) intravenous infusion during daily 6-hr sessions, conducted 5 days per week for 4 weeks (**Fig. 1A**). The training context and parameters are described in detail in **fig. S1** as they are important for interpreting dopamine signals. Briefly, upon the insertion of levers (both active and inactive), the training chamber was lit with light above the active lever (light ON). Mice were allowed to move freely in the chamber, where they exhibited typical spontaneous behaviors, such as locomotion, grooming, and rearing. They were trained to press the active lever at a fixed ratio (from FR 1, 2, to FR4) to obtain intravenous infusion of cocaine of fentanyl. Because the training was self-paced, the interval between lever insertion and drug infusion onset varied substantially across trials and animals (from less than 20 seconds to ten minutes, **fig. S1**). Once drug infusion started, each infusion (lasting ∼2.8 seconds) was accompanied by a 19.5-second ON-OFF blinking of the light above the active lever. After this period, the levers were retracted, and the chamber lights were turned off. Following a 20.5-second dark interval, the next trial began with the lights turned on and the levers reinserted.

**Fig. 1.**
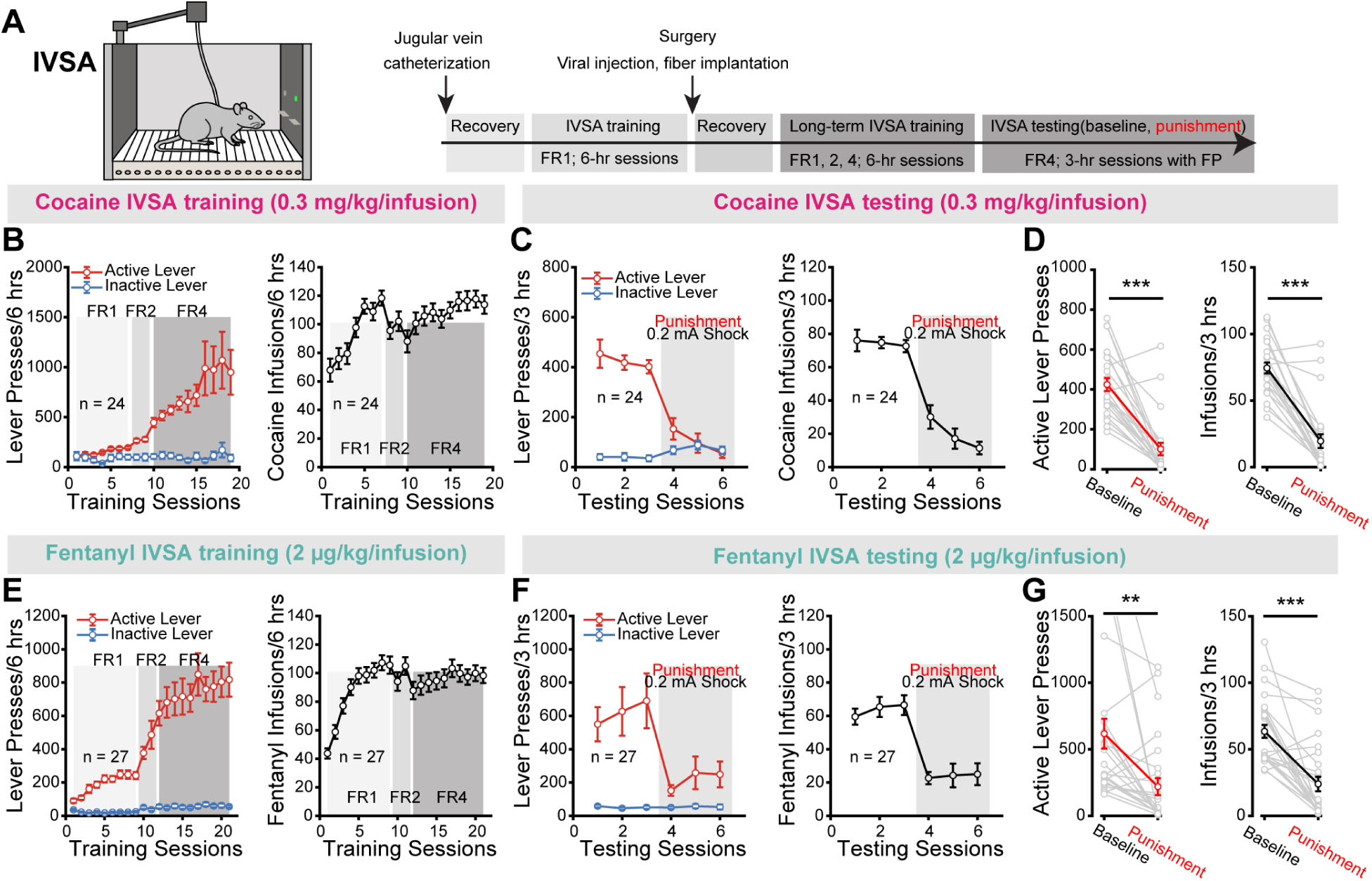
Cocaine and fentanyl intravenous self-administration (IVSA) behaviors. (**A**) Schematic of the IVSA setup and experimental timeline for IVSA training and testing. (**B**) and (**C**) Lever presses (left) and cocaine infusions (right) during the training (**B**) and testing (**C**) phases of cocaine IVSA (n =24 mice). (**D**) Average active lever presses (left) and cocaine infusions (right) during baseline and punishment sessions of the cocaine IVSA testing phase (paired t-test). (**E**-**G**) Same analyses as in **B**-**D**, but for fentanyl (n = 27 mice). Each gray line in (**D**) and (**G**) represents an individual mouse. Red and black lines show group means. Error bars indicate mean ± standard error mean (SEM). ** represents p < 0.01; *** represents p < 0.001.

Consistent with prior studies(*47*–*49*) with long-access drug IVSA, active lever presses and the cocaine (n = 24, **Fig. 1B**) or fentanyl (n = 27, **Fig. 1E**) intakes gradually escalated across the training sessions under the FR1 schedule, while inactive lever presses remained minimal. When the reinforcement schedule was increased to FR2 and subsequently FR4, active lever presses continued to escalate for both cocaine (reaching 997 ± 244 active presses per 6-hour session by the end of FR4 training) and fentanyl (averaging 799 ± 89 active presses per 6-hour session by the end of FR4 training). Drug intake for both cocaine and fentanyl remained high after an initial dip when switched to FR2 (**Fig. 1B**, **1E**). Note that under the FR4 schedule, each lever press had a low probability of resulting in drug infusion. Notably, mice exhibited substantial variability in the latency to the first active lever press upon lever insertion, inter-press intervals, and the latency from lever insertion to drug infusion in both cocaine and fentanyl IVSA (**fig. S1)**. As controls, we also trained a separate group of mice (n = 8) to self-administer saline under a similar FR1 to FR4 schedule. Interestingly, these mice also pressed the active lever slightly more than the inactive lever for saline infusions (**fig. S2)**. But the discrimination between active and inactive lever presses is significantly lower than that of mice IVSA cocaine and fentanyl (**fig. S2**).

Following the 4-week extended training period, mice underwent at least three 3-hour regular cocaine or fentanyl IVSA sessions under the FR4 schedule (baseline sessions) before they were subjected to three 3-hour punishment sessions (**Fig. 1C**, **1F**). During the punishment sessions, each drug infusion (triggered by the 4^th^ active lever press) was paired with a mild foot shock (0.2 mA, 0.5 second). As expected, punishment significantly reduced both the average number of active lever presses and the average intake of cocaine and fentanyl (**Fig. 1D, 1G**). However, there existed clear individual variability in punishment sensitivity with some mice continuing to press the active lever for drug infusions despite receiving foot shocks.

To characterize individual differences in drug-taking and punishment-responsiveness, mice were simply categorized as high or low drug-taking (**Fig. 2A-B** and **2H-I,** orange vs. light blue) and as high or low punishment-resistant (**Fig. 2D-E** and **2K-L,** red vs. dark blue), based on their normalized intake of cocaine or fentanyl during the baseline and punishment sessions, respectively (see **Methods**). Mice whose drug intake fell within mean ±10% of the group mean were unclassified (grey samples in **Fig. 2**). We compared lever-pressing between low- and high-taking groups and observed similar behavioral patterns in the high cocaine- (n = 8) and fentanyl-taking (n = 11) mice. High drug takers showed significantly more active lever presses per infusion than the low drug-takers (**Fig. 2C**, **2J**, 6.3±0.27 vs 5.0±0.08 presses for cocaine; 11.1±1.6 vs 5.8±0.20 presses for fentanyl), indicating that these mice tended to exhibit more futile lever presses (i.e. presses that did not result in additional infusions during the 19.5-second light-blinking period). In addition, high drug-taking mice for both cocaine and fentanyl displayed significantly shorter latencies of the first active lever press following lever insertion (33.6±3.85 vs 105.8±7.75 seconds for cocaine; 39.3±4.45 vs 122.7±12.74 seconds for fentanyl) and shorter inter-press intervals between active lever presses (9.8±0.82 vs 17.9±2.6 seconds for cocaine; 7.9±1.13 vs 23.7±1.69 seconds for fentanyl) compared with the low drug-taking groups. These results indicate that high drug-takers tended to respond more rapidly and persistently to the active lever (**Fig. 2C**, **2J**, also see representative examples in **fig. S1**).

**Fig. 2.**
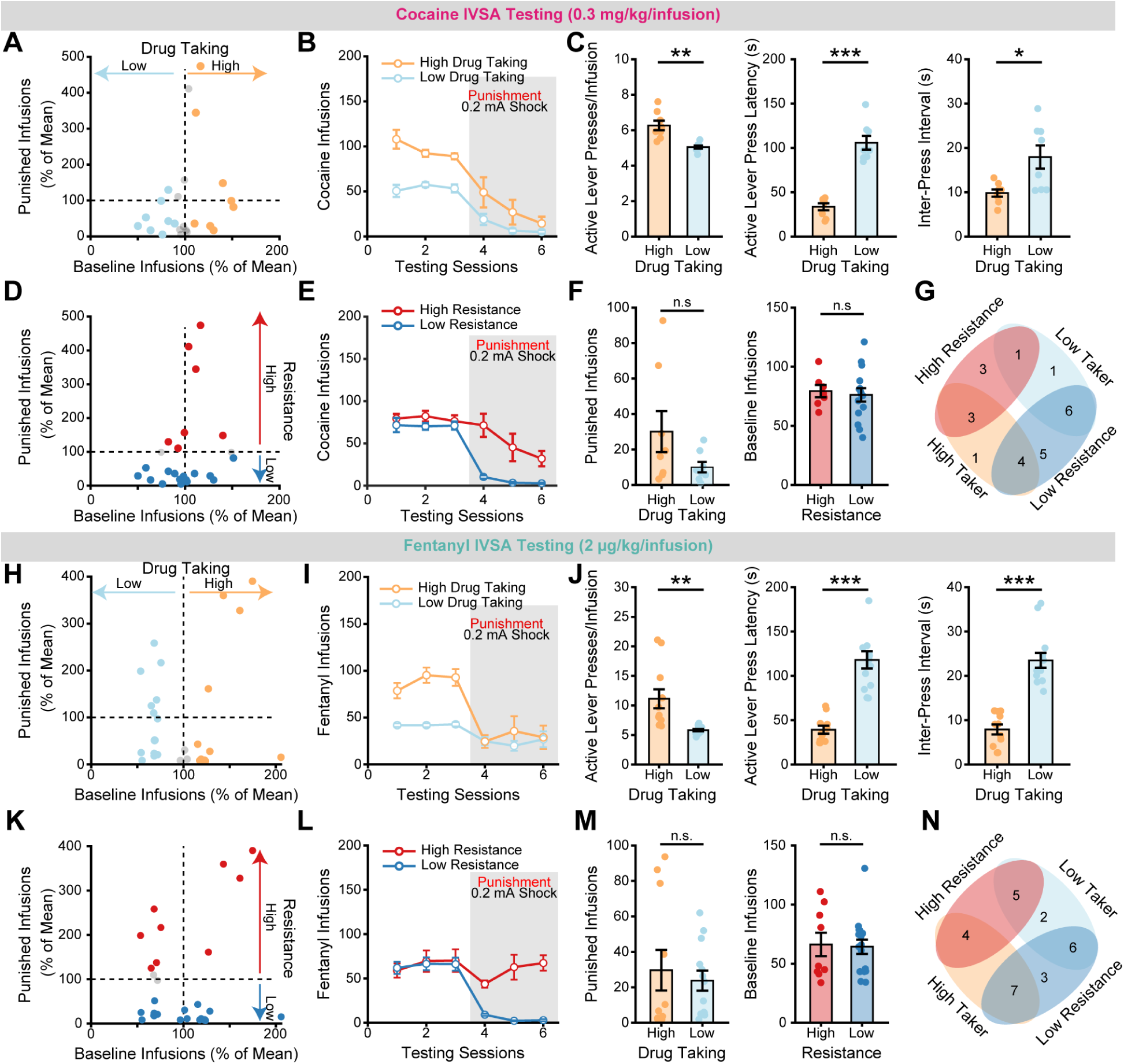
Individual differences in drug taking and punishment resistance. (**A**) Normalized distributions of cocaine infusions (i.e., normalized to the mean) during baseline and punishment. Mice were classified as high (cocaine: n = 8) or low (cocaine: n = 8) drug taking based on whether their baseline infusions were above (orange) or below (light blue) the mean by 10%. (**B**) Cocaine infusions during the testing sessions, grouped by high vs. low drug taking. (**C**) Measures of active lever presses per infusion (left), latency for active lever press to the lever insertion (middle), and inter-press interval (right) during the baseline cocaine IVSA (Welch’s t-test). (**D**) Similar scatter plots as in panel (**A**). Mice were classified as high (cocaine: n = 7) or low (cocaine: n = 15) punishment-resistant based on whether their punished cocaine infusions were above (red) or below (dark blue) the mean by 10%. (**E**) Cocaine infusions during the testing sessions, grouped by high vs. low punishment resistance. (**F**) Cocaine infusions during punishment (left) and baseline (right) sessions, grouped by drug-taking (left) or punishment-resistance (right) categories (Welch’s t-test). (**G**) Overlap between high/low drug-taking and high/low punishment-resistant groups for cocaine IVSA. (**H-N**), Same analyses as in (**A-G**), but for fentanyl. Error bars indicate mean ± standard error mean (SEM). ** represents p < 0.01; *** represents p < 0.001; n.s. represents not significant.

Regarding punishment resistance (**Fig. 2D-E** and **2K-L**), we observed opposite trends of punished drug intake between cocaine- and fentanyl-taking group. High punishment-resistant mice for cocaine (n = 7) tended to decrease their drug intake across the three punishment sessions, whereas high punishment-resistant mice for fentanyl (n = 9) tended to increase their intake over the same punishment sessions (**Fig. 2E**, **2L**), suggesting fentanyl was more effective at promoting punishment-resistance. We next examined whether individuals exhibited correlations between their baseline and punished drug intake. On average, high drug-takers were not significantly more resistant to punishment than low drug-takers under either the cocaine or fentanyl conditions (**Fig. 2F**, **2M**). Likewise, high and low punishment-resistant mice exhibited comparable baseline drug intake (**Fig. 2F**, **2M).** Indeed, there were mixed overlaps between the high vs. low drug-taking and high vs. low punishment-resistant groups (**Fig. 2G**, **2N**). Thus, within the current experimental paradigm, punishment resistance does not correlate with baseline level of drug taking, suggesting different underlying mechanisms modulating these two phenomena.

### Dopamine signatures of excessive drug-taking behavior

Dopamine (DA) plays a crucial role in the reinforcing effects of both cocaine and opioids. But how DA signals are related to individual differences in drug-taking behaviors remain obscure. To examine DA dynamics in the cocaine- and fentanyl-IVSA mice described above, we expressed a genetically encoded DA sensor (AAV2/5-hSyn-GRAB_DA2m)(*50*, *51*) in the dorsal medial shell of the NAc and implanted an optic fiber to collect fluorescent signals (**fig. S3, Fig. 3A** and **3J**). Using rotary fiber photometry (FP), which is compatible with the IVSA in freely behaving mice, we monitored the DA dynamics during cocaine and fentanyl IVSA in fully trained mice across 3-hour baseline testing sessions under the FR4 schedule (timeline shown in **Fig. 1A**). To minimize photobleaching from continuous long-duration recording, we performed three cycles of 30 minutes rotary FP, each separated by a 30-minute interval (i.e., three episodes of 30-minute recording per session), and concatenated them for analysis.

**Fig. 3.**
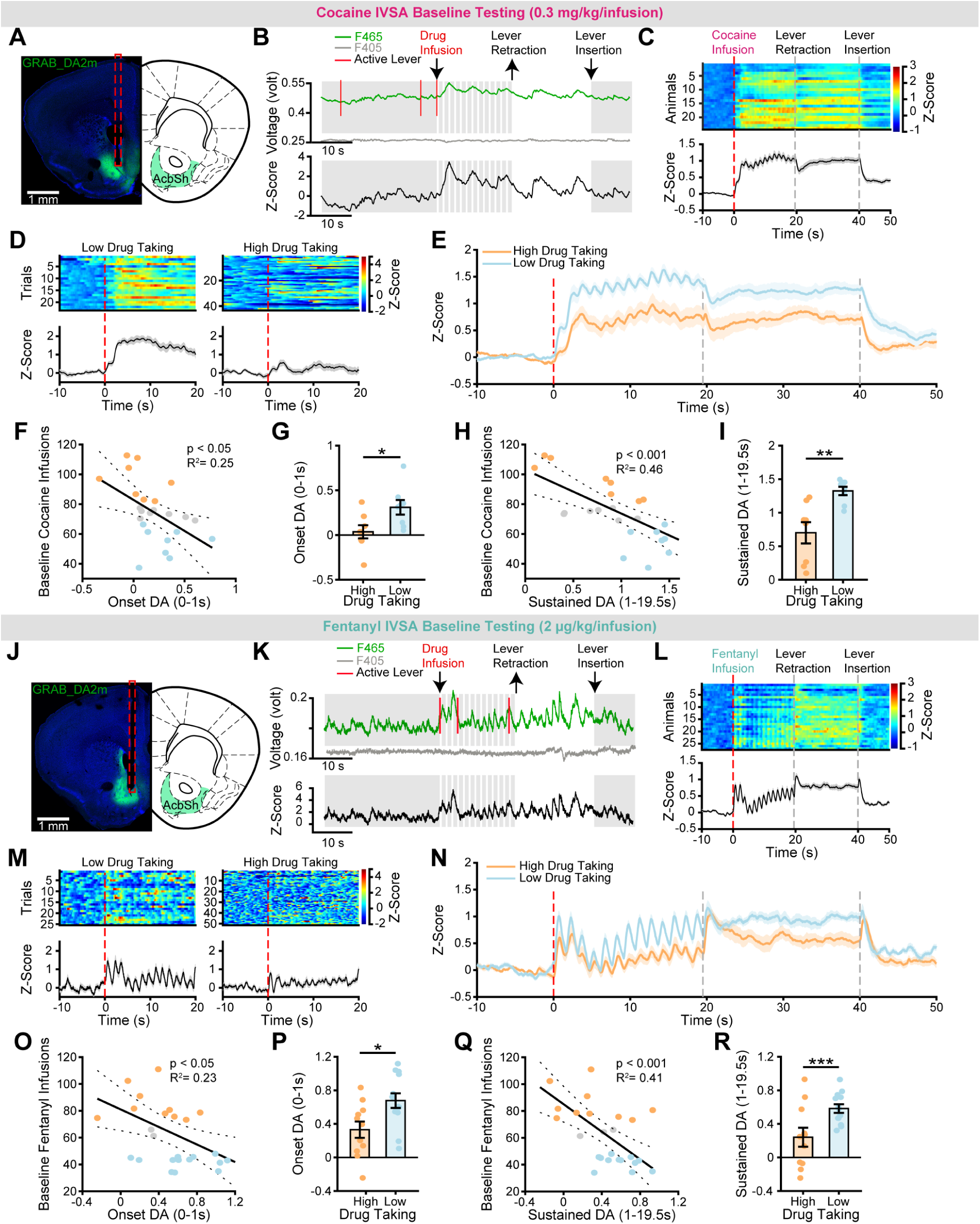
Dopamine dynamics during cocaine and fentanyl self-administration after extended training. (**A**) Images showing GRAB_DA2m (green) expression and fiber tracks (red dashed line) in the NAc medial shell from a cocaine-IVSA example. (**B**) Top: example raw photometry traces (465 nm [green] and 405 nm [gray]) with behavioral events overlaid during cocaine IVSA. Bottom: z-score of DA signals after preprocessing the raw data; gray shading indicates cue light on/off. (**C**) Group DA responses to contingent cocaine infusions (n = 24). Each row of the colormap represents one mouse, sorted by the number of infusions (top = most). The red dash lines at time 0 represent the start of drug infusion. The gray dash lines represent lever retraction and insertion. (**D**) Example DA responses to contingent cocaine infusions from a low (left) and a high (right) drug-taking mice. (**E**) Time courses of DA responses to contingent cocaine infusions for high (orange) and low (light blue) drug-taking mice. (**F**) Linear regression showing the relationship between onset DA responses and number of baseline infusions of cocaine. (**G**) Bar graphs quantifying onset DA responses to cocaine infusions in high vs. low drug-taking mice (Welch’s t-test). (**H**) Linear regression showing the relationship between sustained DA responses and number of baseline infusions of cocaine. (**I**) Bar graphs quantifying sustained DA responses to cocaine infusions in high vs. low drug-taking mice (Welch’s t-test). Orange dots represent high drug-taking mice, light blue dots represent low drug-taking mice and gray dots represent mice not classified. (**J-R**) Same analyses as in (**A-I**), but for fentanyl (n = 27). Error bars indicate mean ± standard error mean (SEM). * represents p < 0.05; ** represents p < 0.01; *** represents p < 0.001

Examining the raw and z-scored FP traces across different trial epochs (see representative examples in **Fig. 3B** and **3K**), we found that in both the cocaine and fentanyl IVSA, DA sensor signals were relatively low prior to drug infusion. At the onset of infusion, DA signals increased, with multiple peaks (referred to as DA transients) that occurred during the drug infusion, the light-blinking period, and the lever-retraction/light OFF (inter-trial interval) phases. Once the light was turned back ON and the levers were reinserted to initiate the next trial, DA signals declined. Notably, the width of individual DA transients was wider in cocaine-IVSA compared to fentanyl-IVSA conditions. This is likely due to cocaine’s blockade of the DA reuptake transporter, which slows the clearance of extracellular DA and results in a significantly slower decay of DA signals (Decay slope: -0.54±0.04 vs -1.28±0.05, **fig. S4**). By contrast, no consistent DA signals were observed at the time of active lever press (**fig. S5**), likely reflecting the fact that at FR4, each lever press has a low probability of resulting in drug reward (also see computational modeling below).

We computed the time course of z-scored DA signals for each animal (see **Methods**) and plotted averaged z-scores of all trials aligned to the onset of drug infusion, spanning from 10 seconds before infusion to 10 seconds after the initiation of the next trial (n=24 mice for cocaine; n=27 mice for fentanyl, **Fig. 3C**, **3L**). During cocaine IVSA, the averaged DA levels increased at the onset of infusion and remained elevated throughout the light-blinking cue period and the inter-trial interval (lights OFF period) (**Fig. 3C**). Similarly, in mice during fentanyl IVSA, increased DA levels also occurred at the onset of infusion, and the DA elevation displayed phasic, oscillatory responses during the light-blinking period, which then became a sustained elevation during the 20.5-second dark inter-trial interval (**Fig. 3L**). The oscillatory DA pattern was more clearly observed in the group-averaged signals for both fentanyl-IVSA and cocaine-IVSA during the light-blinking period (lower panels in **Fig. 3C**, **3L**). Notably, the rise-and-fall DA signals peak shortly after each OFF-ON transition of the blinking light (**fig. S3D**), indicating the averaged oscillations reflected responses to the light cue. In both cocaine and fentanyl IVSA, DA levels dropped rapidly at the onset of the next trial when the light turned ON and levers were inserted (**Fig. 3C and 3L**). Importantly, the drug-infusion and blinking-light evoked DA responses were absent on the first day of training (e.g., the auto-shaping session) and in saline-IVSA mice (**fig. S3E-F**), indicating that these signals emerged through drug-cue associative learning. In contrast, the elevated DA signals during the light OFF inter-trial interval were also present on the first day of training and in saline-IVSA mice, possibly reflecting mice’ natural preference for darkness.

Given these observations, we focused our analyses on the infusion and light-blinking periods for comparing DA dynamics between high and low drug-taking groups. Representative DA signals of individual trials from a low- and high-cocaine taker (**Fig. 3D**), along with group-averaged signals (**Fig. 3E**) revealed interesting differences. While DA responses varied from trial to trial, differing in both amplitude and the timing of peak z-score, critically, low cocaine-taking mice consistently showed larger DA increases than high takers (**Fig. 3D-E**). This result is reminiscent of previous studies showing that rats exhibiting escalated cocaine intake across training displayed reduced DA to contingent cocaine infusion(*27*, *28*). Similarly, low-fentanyl taking mice also consistently exhibited larger DA increases than high-fentanyl takers, despite large trial-to-trial variations in DA signals (**Fig. 3M-N**). We separately quantified the DA responses at the infusion onset (0-1 second from infusion start, referred to as “onset DA signal”) and during the subsequent cue period (1-19.5 second from infusion start, referred to as sustained DA signal). Importantly, linear regression analysis revealed that both the onset and sustained DA signals were significantly negatively correlated with the number of cocaine (**Fig. 3F** and **3H**) or fentanyl (**Fig. 3O** and **3Q**) self-administrations. Group comparison further confirmed that high drug-taking mice showed significantly smaller onset (**Fig. 3G** and **3P**) and sustained DA signals (**Fig. 3I** and **3R**). The reduced DA signals in high drug-takers are unlikely to be caused by elevated baseline DA levels that might blunt additional drug-evoked responses. If this were the case, DA responses should decrease over the course of a session as more drug is consumed. However, when we compared DA responses across the three recording episodes (the 1st, 2nd, and 3rd 30-minute blocks), we found no evidence of a progressive decline in evoked DA (**fig. S6**).

Altogether, despite the distinct DA dynamics evoked by cocaine and fentanyl IVSA, both drugs showed a consistent relationship between drug-taking behavior and DA responses: higher drug intake was associated with weaker DA responses to contingent drug infusion and drug-associated cues.

### Dopamine signatures of punishment resistance

Compulsive drug-taking despite adverse consequences is a hallmark of drug addiction, thus we examined DA responses in the NAc medial shell during punishment sessions. Following the co-occurrence of drug infusion with a mild foot shock (0.2 mA, 0.5 s), average DA signals in both the cocaine and fentanyl groups showed a marked increased during the post-shock/infusion period and remained elevated throughout the light-blinking, and inter-trial dark periods (**Fig. 4A, 4I**). Overall, cocaine infusion plus shock produced variable onset DA responses (0-1 second post-infusion), followed by a uniform sustained DA surge (1-19.5 second post-infusion) (**Fig. 4A**). Closer examination of individual mice in the cocaine group revealed heterogeneity in DA dynamics: in a subset of mice, the co-occurrence of cocaine infusion and shock elicited an initial dip or pause in DA followed by a robust, sustained rebound increase of DA, whereas in others, the infusion and shock triggered an instant increase in DA that remained elevated (**fig. S7, Fig. 4B**). Visual inspection of DA dynamics in high and low punishment-resistant mice revealed that both groups had comparable onset DA responses, but the low-resistant mice exhibited significantly larger sustained DA signals (**Fig. 4C-D**). Quantitative analyses confirmed that group-averaged infusion/shock-evoked onset DA was neither significantly different between high- and low-resistant mice, nor correlated with the amount of punished cocaine intake (**Fig. 4E-F**). However, at the individual level, 40% (6/15) of low-resistant mice displayed a suppression of onset DA, compared to only 14.3% (1/7) of high-resistant mice (**Fig. 4F**, Fisher’s exact test, p = 0.35). In contrast, there is a strong and significant negative correlation between the post-shock sustained DA levels and the punished cocaine infusions: high-punishment resistant mice had low sustained DA levels (**Fig. 4G-H**). As a control, co-occurrence of saline infusion and shock predominantly elicited a dip in onset DA responses (83.3%, [5/6]), followed by a rebound (**fig. S3G**).

**Fig. 4.**
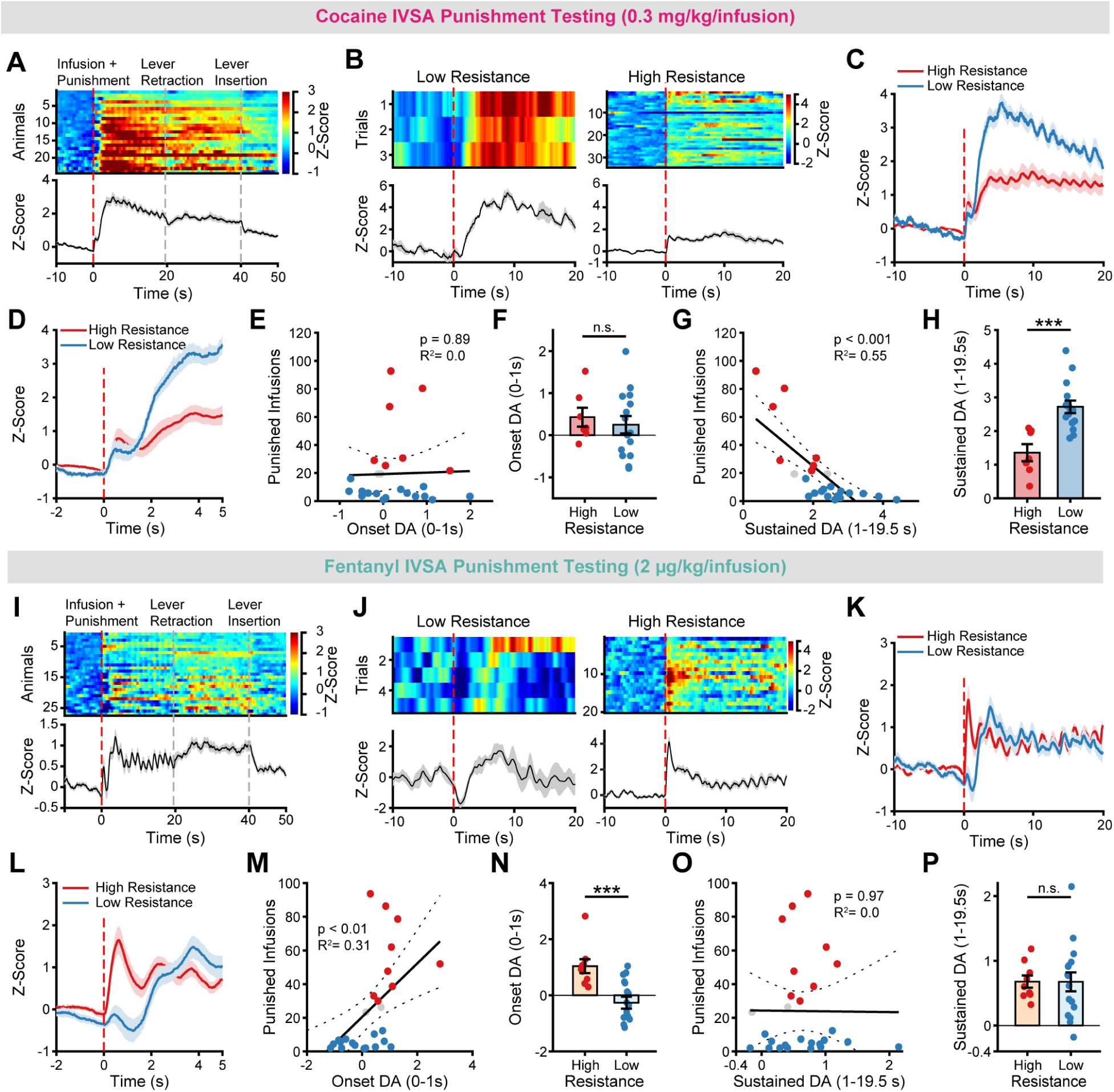
Dopamine dynamics during punished drug taking. (**A**) Group DA responses to punished cocaine infusions (n = 24 mice). Each row of the colormap represents one mouse, sorted by the number of punished infusions (top = most). The red dash lines at time 0 represent the co-occurrence of drug IVSA and punishment. The gray dash lines represent lever retraction and insertion. (**B**) Example DA responses to punished cocaine infusions from a low (left) and a high punishment-resistant (right) mice. (**C**) Time courses of DA responses to punished cocaine infusions for high- (red) and low-resistant (blue) mice. (**D**) Same as (**C**), but zoomed in to highlight the onset response (0-1 s post-infusion). (**E**) Linear regression showing the relationship between onset DA responses and number of punished infusions of cocaine. (**F**) Bar graphs quantifying onset DA responses to punished cocaine infusions in high- vs. low-resistant mice. (**G**) Linear regression showing the relationship between sustained DA responses and the number of punished infusions of cocaine. (**H**) Bar graphs quantifying sustained DA responses to punished cocaine infusions in high- vs. low-resistant mice. Red dots represent punishment-resistant mice, blue dots represent punishment-sensitive mice and grey dots represent mice not classified. (**I-P**) Same analyses as in (**A-H**), but for fentanyl. Error bars indicate mean ± standard error mean (SEM). *** represents p < 0.001; n.s. represents not significant.

In the fentanyl group, the co-occurrence of fentanyl infusion with shock also elicited heterogeneous onset DA responses across mice (**fig. S6, Fig. 4I, 4J**). Visual inspection of the group-averaged DA dynamics showed that high punishment-resistant mice exhibited a sharp increase in onset DA, whereas the low-resistant group displayed a dip of onset DA signal; however, both groups showed comparable sustained DA (**Fig. 4J-L**). Quantitative analyses confirmed that the fentanyl-infusion/shock-evoked onset DA response was significantly higher in high-resistant mice and was significantly positively correlated with punished fentanyl infusions (**Fig. 4M-N**). Moreover, 56% (9/16) of low-resistant mice showed a reduction in onset DA while none (0/9) of the high-resistance exhibited such a dip (**Fig. 4N**, Fisher’s exact test, p < 0.01). In contrast, the post-shock sustained DA signals were neither significantly different between high- and low-resistant mice, nor correlated with punished fentanyl intake (**Fig. 4O-P**). Among the nine punishment-resistant mice, we also recorded DA responses during subsequent punishment sessions in 8 mice. At the group level, these mice showed a trend toward reduced sustained DA signaling during subsequent punishment sessions, and five of eight exhibited a further increase in onset DA responses (**fig. S8**).

Taken together, in cocaine self-administering mice, the sustained DA responses after shock negatively correlated with resistance to punishment, whereas in fentanyl self-administering mice, the onset DA responses to shock/drug-infusion positively correlated with resistance to punishment.

### A computational model captures DA dynamics in the IVSA paradigm across drugs and conditions

A long-standing theory for NAc DA activity posits that phasic DA signals encode the temporal-difference reward prediction error (TD-RPE), i.e. the mismatch between the expected values of temporally adjacent states(*52*–*54*). In the fentanyl group, punishment-resistant mice displayed a sharp increase in DA at the onset of shock and fentanyl infusion (**Fig. 4L**), a pattern reminiscent of a positive RPE signal observed in Pavlovian conditioning. However, it is unclear whether the existing TD-RPE model of DA signals based on short timescale classical conditioning could model the substantial variabilities in IVSA behaviors (**fig. S1**) and explain the highly heterogenous onset and sustained DA dynamics during both cocaine and fentanyl IVSA and punishment sessions (**Fig. 3** and **Fig. 4**). To address these questions, we modeled each animal as an *Actor-Critic* TD-learning agent performing a self-paced drug self-administration task (**Fig. 5-6**). One key aspect of our model is to represent the distinct epochs within IVSA trials as *internal states, S*(*t*). The agent learns the state value, *V*(*S*(*t*)), and action value, *M*_*a*,*S*(*t*)_, to maximize the expected total future reward using the TD-RPE signal, δ(t). Thus, despite the low predictivity of individual lever presses for drug reward at FR4, the agent learns that pressing the lever is the best policy (**see Methods**). We used model-derived δ(t) to generate δ̃(t) as the simulated DA signals by clipping the negative value to -0.5 and adding a decay tail to δ(t). We then compared δ̃(t) with the experimentally observed NAc DA dynamics and analyzed the correlations between δ̃(t) and drug taking or punishment resistance.

**Fig. 5.**
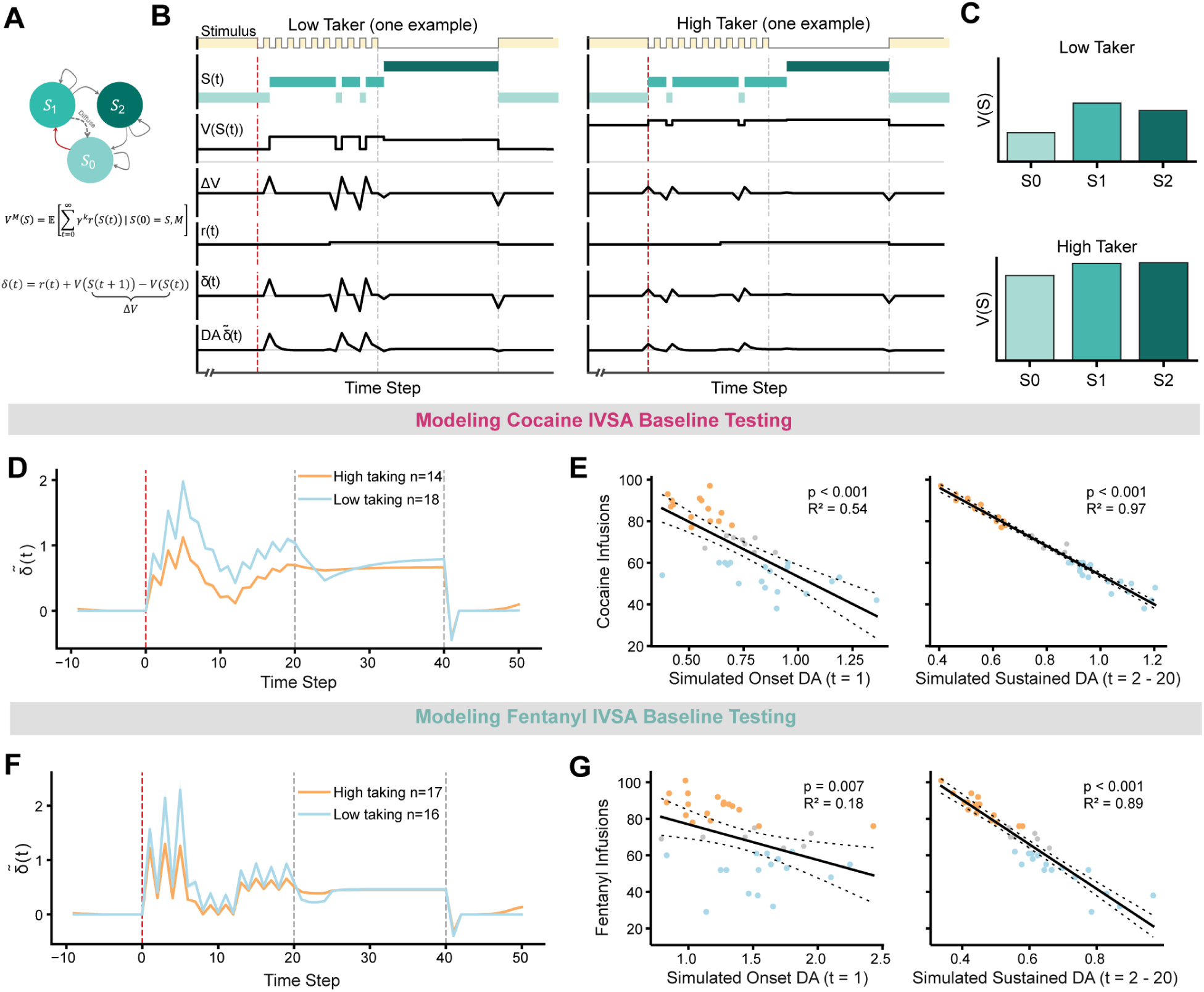
Actor-Critic TD learning model can explain DA dynamics and negative correlation between DA signals with drug intake during drug IVSA tasks. (**A**) Schematic of the model, highlighting the three discrete internal states (S0, S1, S2) and their transition. (**B**) Simulation results showing the internal state, state value, change of value of temporally adjacent states, drug reward, TD error δ(t) and simulated DA δ̃(t), at each timestep in simulated trials for a low-taking (left) and high-taking agent (right). (**C**) State value of example low-taking (top) and high-taking agents (bottom). (**D**) Average δ̃(t) signals for high and low cocaine-taking agents. (**E**) Correlation between onset and sustained δ̃(t) with simulated cocaine infusions. (**F-G**) Same analyses as in (**D-E**), but for fentanyl IVSA agents.

**Fig. 6.**
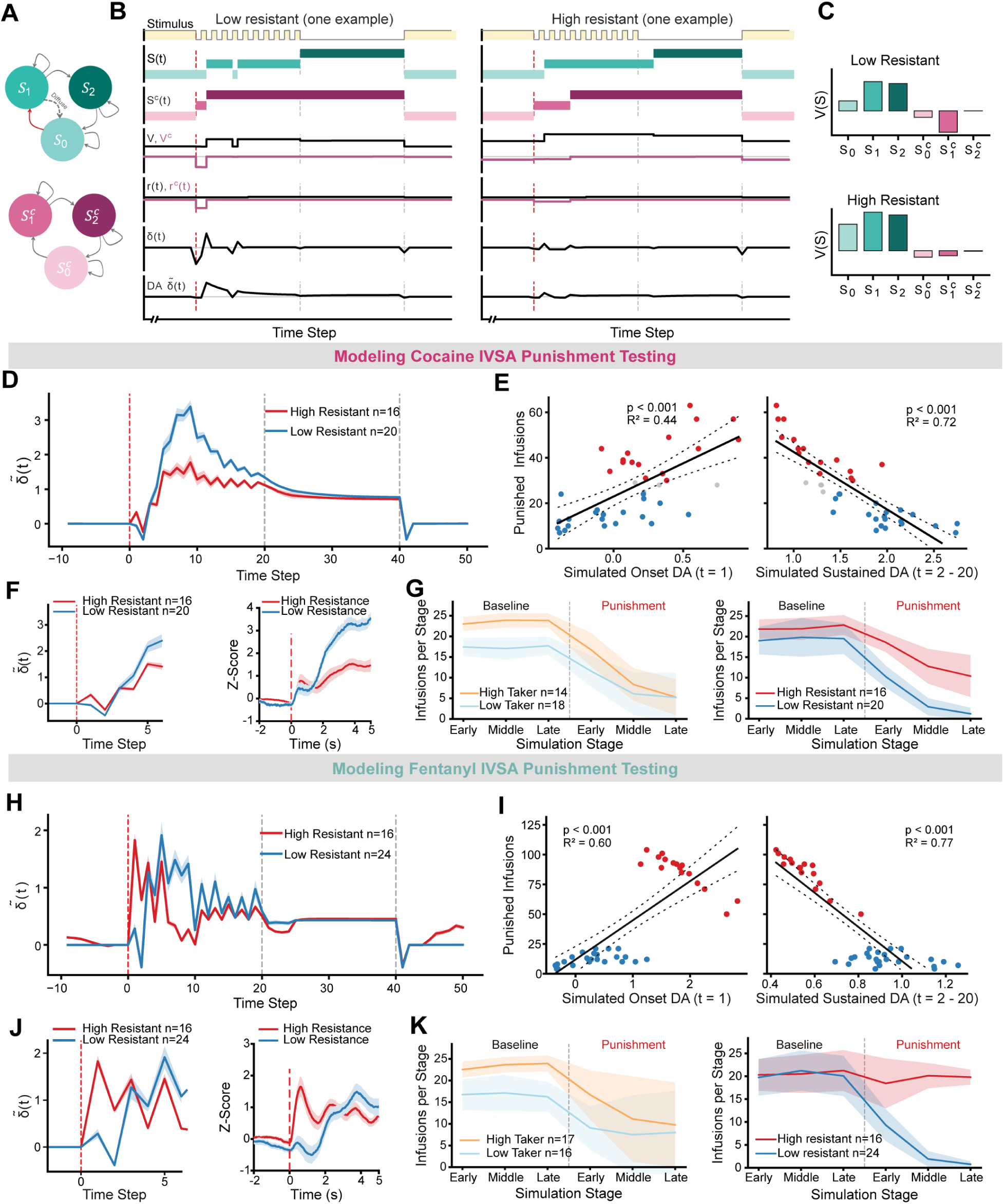
Actor-Critic TD learning model can partially explain DA dynamics and the correlation between DA signals with punishment resistance during punishment sessions of IVSA tasks. (**A**) Schematic of the model, highlighting the three discrete internal states (S0, S1, S2), shock-modulated states (S^C^0, S^C^1, S^C^2) and their transition. (**B**) Simulation results showing the state, state value, change of value of temporally adjacent states, drug reward, shock reward, TD error δ(t), and simulated DA δ̃(t), at each timestep in simulated cocaine trials for a low-resistant (left) and high-resistant agent (right) during a punishment session. (**C**) State value of example low-resistant (top) and high-resistant agents (bottom). (**D**) Average δ̃(t) signals for high and low punishment-resistant cocaine-taking agents. (**E**) Correlation between simulated onset and sustained DA with simulated punished cocaine infusions. (**F**) Left panel, same as (**D**), but zoomed in to highlight the onset response. Right panel, experimental data as showed in **Fig. 4D**. (**G**) The count of cocaine infusions by agents during baseline and punishment sessions across simulation stages, grouped by high and low drug taking (left) or punishment resistance (right). (**H-K**), same analyses as in (**D-G**), but for fentanyl taking agents.

We first modeled the FR4 drug-taking sessions. Each trial was modeled as inducing three discrete internal states, S_0_, S_1_ and S_2_, defined by their distinct sensory contexts (**Fig. 5**): S_0_ denotes the period from lever insertion and light ON to the moment of drug infusion—a phase that typically occupies most of the trial and can last from seconds to minutes. S_1_ denotes the drug infusion and associated light-blinking period, whereas S_2_ corresponds to the dark inter-trial interval (ITI). For simplicity, the external sensory stimuli associated with S_1_ and S_2_ were each modeled as lasting 20 timesteps (t). We further assumed that a drug reward r_d_ becomes perceptible 10 timesteps after drug infusion and persists to the end of the trial. A critical novel aspect of the model is to incorporate uncertainty in the agent’s estimation of current states S(t). For instance, the agent may fail to notice the first few blinks of the light cue, leading to temporal variation in its estimate of when the light-blink state begins, or may confuse states that share similar sensory features (e.g. a blink-ON moment in S_1_may resemble the light-on period in S_0_).

During self-administration, the learned value of state *S*_1_, *V*(*S*_1_), is approximately constant across individual agents since *S*_1_ has a fixed reward contingency by task design. In contrast, state *S*_0_ acquired predictive value from drug infusions in preceding trials, with different agents learning different *V*(*S*_0_). Critically, the strength of the *S*_0_ → *S*_1_contingency (red arrow in **Fig. 5A**) depends on the learned action value *M*_*a*,*S*_0__ which determines both the probability and timescale of lever presses (**see Methods**). High drug takers, which select the lever press action more frequently with shorter inter-press intervals, learn that sufficient lever presses in *S*_0_ reliably lead to *S*_1_ and drug reward. As a result, they acquire a high expected *V*(*S*_0_) that approaches *V*(*S*_1_), yielding a small prediction error at the *S*_0_ → *S*_1_ transition (because *δ*_{0→1}_ ≡ *V*(*S*_1_) − *V*(*S*_0_)). In contrast, low drug takers are less certain that lever pressing drives the *S*_0_ → *S*_1_ transition. Therefore, compared to high takers, they select the lever press action less frequently, with longer inter-press intervals, and maintain *V*(*S*_0_) ≪ *V*(*S*_1_), resulting in a larger *δ*_{0→1}_ upon entry into *S*_1_. Altogether, *δ*_{0→1}_ at the onset of drug infusion (timestep 1 in *S*_1_) is negatively correlated with the amount of drug taking (**Fig. 5 B-C**).

Importantly, *δ*_{0→1}_ also contributes to *δ*(t) during the light-blinking period of *S*_1_ due to animals’ uncertainty of state transitions. Attentional lapse (modeled as a ≈ 6 timestep jitter) in detecting the drug-associated cue (light blinking off) generates multiple cue-locked *δ*(t) peaks at the initial phase of *S*_1_. In addition, *S*_1_ → *S*_0_ confusion (light blinking on) further produces cue-locked oscillations in δ(t) when averaged across trials (**Fig. 5F, see Methods**). Moreover, the delayed perception of drug reward, *r*_*d*_(*t*) induces a slow rise of *δ*(t) starting ≈ 10 timesteps after infusion onset, which is superimposed on the oscillation of *δ*(t). Taken together, the average sustained δ(t) or δ̃(t) over the post-infusion phase of *S*_1_ (timesteps 2-20), is determined by *δ*_{0→1}_ and *r*_*d*_(*t*), and therefore is also negatively correlated with drug intake (**Fig. 5G**). Finally, we assumed that the effect of the drug reward extends into *S*_2_. Upon the *S*_2_ → *S*_0_ transition, although light on and lever insertion predict the next reward cycle, termination of the prolonged drug reward mainly produces a negative *δ*_{2→0}_.

The simulated DA *δ̃(t)* traces of FR4 drug-taking closely resemble the actual DA signals observed in the fentanyl IVSA experiments and recapitulate the negative correlations between both onset and sustained DA signals with fentanyl intake (**Fig. 5F-G**). When the decay time constant of δ̃(t) is increased fivefold to mimic cocaine-caused inhibition of DA reuptake, the exponentially filtered δ̃(t) traces similarly reproduce the DA dynamics observed during cocaine IVSA, as well as the negative correlations between DA signals and cocaine intake (**Fig. 5D-E**). Together, these results indicate that despite the complexity and temporally extended nature of the IVSA paradigm and the distinct pharmacological classes of the drugs (psychostimulant vs. opioid), contingent DA release in the NAc during these tasks can be uniformly explained as encoding TD-RPE.

Using the same Actor-Critic TD-learning framework, we next modeled the punishment sessions. We assumed that footshock induces an additional internal state cascade *S*^*c*^(*t*) (**Fig. 6**), which runs in parallel with the three task-related states 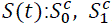, *and* 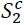. Here, 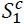 denotes the immediate shock-alert state, and the 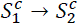 transition occurs stochastically as the agent settles into a shock-relieved “safety” state 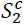. At the onset of the next trial (light on and lever insertion), the shock-related state returns to a baseline “danger” state 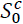, which persists until drug infusion and the next shock. Accordingly, the agent undergoes a transition from 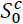 *to* 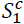 at the drug infusion/shock onset, followed by a stochastic transition 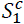 *to* 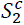, corresponding to shock relief. In the experiments, a footshock is delivered at the 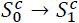 transition to induce a negative reward *r*_*c*_ < 0. In addition, we initialized 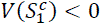 to reflect the innate aversive valuation of the shock-alert state. Given the task states *S*(*t*) and shock states *S*^*c*^(*t*), we assume additive state and action-value components:

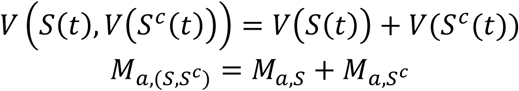

We modeled fentanyl and cocaine as having different effect on changing a threshold *θ*_*c*_ in the perception of shock stimuli as aversive. Accordingly, if the delivered shock level falls below *θ*_*c*_ it contributes to sensory salience; while if it falls above *θ*_*c*_, it contributes to negative valence. Thus, *θ*_*c*_ modulates the agent’s learning rate (**see Methods**).

During cocaine IVSA with punishment, 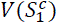 is negative at the onset of drug infusion (**Fig. 6B-C**). Spontaneous 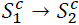 transitions during the post-shock period produce a large delayed δ(t) surge due to 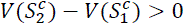. Together with the δ(t) derived from the delayed drug reward, the total δ̃(t) (that considered cocaine-induced slow decay) exhibits sustained increase during the post-shock light-blinking period. This pattern closely recapitulates the large and sustained DA signals observed experimentally (**Fig. 6D**). Because the shock-relief component of this sustained δ̃(t) reflects agents’ shock sensitivity, it is negatively correlated with punishment resistance: individuals with low resistance exhibit more sustained δ̃(t), consistent with the experimental findings (**Fig. 6D-E**).

During fentanyl IVSA with punishment, although 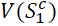 is negative in the early trials, in punishment-resistant agents, we model fentanyl to progressively suppresses shock sensitivity (increases the threshold *θ*_*c*_). As a result, the initially aversive unconditional stimulus (US, footshock) gradually becomes a salient conditioned stimulus (CS+) predictive of fentanyl reward in these individuals. Thus, the model generates a strong positive δ(t) or δ̃(t) transient at the 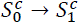 transition in punishment-resistant fentanyl-taking agents but not in cocaine-taking agents (**Fig. 6J, 6F**), matching the different DA signals from the two drugs in experiments (**Fig. 4D, L**). This reversal of US to CS+ does not occur in punishment sensitive individuals (**Fig. 6J**). These δ̃(t) dynamics recapitulate the positive correlation between onset DA responses and punishment-resistant fentanyl taking in experiments (**Fig. 6H-I**). Moreover, we plotted the drug infusion numbers for high- vs low-taking agents and high- vs low-punishment resistance agents from baseline FR4 sessions to the three consecutive punishment sessions (**Fig. 6G** and **6K**), and found that agents in the model also mimic the drug-taking patterns observed in mice (**Fig. 2B**, **2E**, **2I**, **2L**).

Our model also predicted a weaker but significant positive correlation of onset δ̃(t) with punished cocaine intake, as well as negative but significant correlation of sustained δ̃(t) with punished fentanyl intake, both of which were not clearly observed experimentally. These discrepancies may reflect the complexity of how different drugs modulate the computation of value and RPE that is not considered by the model. Nevertheless, overall, the computational model captures the diversity of drug-taking and punishment-responsiveness behaviors, along with the associated DA dynamics across trials epochs, individuals, and drug classes. Importantly, it reveals a simple, unified role of NAc DA signals in encoding TD-RPE across different phases of addiction.

## Discussion

Here, we used a genetically encoded DA sensor to characterize the dynamic patterns of DA release in the NAc medial shell in a mouse model of drug addiction, comparing DA responses to cocaine and fentanyl IVSA in high versus low drug takers, as well as in animals exhibiting high versus low punishment resistance (i.e., a model of compulsive drug use despite adverse consequences). We found that after extended training under an FR4 schedule, contingent cocaine infusions evoked a sustained increase in DA during the drug-associated cue period (i.e., blinking lights), whereas contingent fentanyl infusions elicited a large increase of DA at infusion onset that evolved into oscillations synchronized with the blinking lights. Similar oscillatory DA patterns could also be observed during cocaine IVSA after responses were averaged across trials and mice, although with smaller amplitudes. Critically, across both cocaine and fentanyl, individual’s drug intake was consistently negatively correlated with DA responses: higher levels of drug intake were associated with lower evoked DA signals. With regards to punished drug taking, we observed distinct DA signatures associated with compulsivity in cocaine versus fentanyl taking. In cocaine IVSA mice, higher punishment resistance was associated with weaker sustained DA responses during the drug-associated cue period, whereas in fentanyl IVSA mice, higher punishment resistance was associated with stronger onset DA responses at the co-occurrence of punishment and drug infusion. To account for these diverse DA dynamics across baseline and punishment sessions, across individuals and across drug classes, we developed an Actor-Critic TD-learning based computational framework that incorporates internal states, agent’s uncertainty, and drug-specific effects. This model captures the observed behavioral diversity and supports a unified interpretation of NAc DA responses as encoding temporal-difference reward prediction errors (TD-RPE) based on internal state estimation.

Many previous computational models of addiction (e.g. opponent theory, incentive salience sensitization theory, habit formation) focused on reproducing addiction behaviors instead of accounting for the DA dynamics as a learning outcome. Here our actor-critic model considers both behavior and DA dynamics. However, different from traditional TD-RPE models of classical conditioning, our framework operates on a self-administration task-relevant timescale and explicitly incorporates both states and action values. We modeled how agents evaluate their states after learning the self-administration task at FR4. Our results reveal that the difficulty of the FR4 task schedule introduces significant sources of uncertainty that naturally give rise to complex dynamics in the RPE signal which resemble the observed DA activity. A previous TD-RPE model proposed by Redish and colleagues(*33*, *34*) relied on the assumption of an un-cancellable RPE elicited by drugs upon infusion, which caused unbounded growth of the drug value and consequently would predict a positive correlation between DA signals and drug taking— inconsistent with experimental observations. In contrast, our model naturally recapitulates the negative correlation between DA signals and contingent drug intake. In our model, individual differences in DA response reflect differences in the learned contingency between task states and drug reward.

Could the finding that NAc DA universally encodes TD-RPE help reconcile the divergent views of DA function in addiction? Two influential and seemingly opposing frameworks have been proposed: the DA depletion hypothesis(*55*) and the incentive sensitization theory (IST)(*39*). Preclinical studies have showed that escalation of cocaine self-administration in rodents is accompanied by reduced phasic DA signaling(*27*, *28*, *43*) and human imaging studies also have reported decreased striatal DA responses in individuals with cocaine use disorders(*29*). However, far less was known about DA signaling in opioid self-administration models. Our findings that excessive intake of both cocaine and fentanyl is associated with blunted DA responses to contingent drug infusion, together with prior studies, appear to be consistent with the hypo-dopamine hypothesis. By contrast, the IST theory, also supported by substantial experimental evidence, posits that drug exposure produces a hyper-responsive DA system, leading to exaggerated phasic DA responses to drug-associated cues and context that drive heightened “wanting” to take drugs.

We believe these two theories are not necessarily contradictory and can be unified by the framework that phasic DA encodes TD-RPE, i.e, the difference between the expected values of temporally adjacent states. As TD-learning agents, animals and humans continuously assign expected values to their current states, with these values shaped by experience and learning history. In the IVSA paradigm, high drug takers learn that sufficient lever presses reliably result in drug infusion, motivating more frequent lever press with shorter inter-press intervals. Consequently, they acquire a high expected value for the lever-pressing state and generate relatively small DA/TD-RPE signals upon receipt of the actual drug infusion and its associated cues. In contrast, low drug takers are less certain that lever pressing will result in drug infusion and are therefore less motivated to press the lever, leading to longer inter-press intervals. Consequently, the actual drug infusion is more unexpected, resulting in a large DA/TD-RPE signals. Thus, under contingent conditions, akin to knowingly taking the drug and fully expecting its effect, more excessive drug taking is associated with lower DA responses. However, in humans with SUD, drug-associated cues can be encountered outside the drug-taking context and robustly cause craving. In such situations, drug cues may elicit higher expected value based on remembered drug reward than the perceived value of an individual’s current state, thereby resulting in large DA/TD-PRE signals, consistent with IST. Although we did not directly assess DA responses to unexpected drug-associated cues outside the IVSA context, a recent study showed that DA release is indeed enhanced in response to non-contingent or unexpected cocaine-paired cues, but diminished when the same cues were encountered in a contingent, predictable context(*28*).

Encoding DA as TD-RPE also provides new insights into compulsive drug taking despite punishment. When drug use is associated with adverse consequences, individuals must decide whether to abstain from further drug taking or to endure punishment to continue pursuing the drug. Drug-induced reduced sensitivity to punishment, deficits in inhibitory control or impairments in punishment learning may bias this decision toward compulsive drug taking despite negative consequences, one of the most intractable features of addiction. Using foot shock as a punishment, we showed that in saline control mice, DA signals in the NAc medial shell were predominantly suppressed upon shock delivery (**fig. S4**), consistent with a strong negative valence of shock for drug-naive animals and with DA encoding a negative TD-RPE. Similarly, many punishment-sensitive mice also showed a suppression of the DA signal at the shock-infusion onset during both cocaine and fentanyl IVSA (**Fig. 4E-F, M-N**). In contrast, this shock-elicited DA dip was absent in all but one punishment-resistant animals, suggesting an impairment of encoding negative RPE. A prior study also found that cocaine exposure disrupted the pause firing of DA neurons in response to reward omission(*56*). We also observed a large-amplitude rebound in DA following shock, which we interpreted as positive RPE reflecting “shock relief”. The rebound signal was present in punished saline- and cocaine-IVSA mice, as well as in punishment-sensitive fentanyl-taking mice. It is plausible that stronger relief signals may provide more robust negative feedback, reinforcing avoidance of the punished action. Consistent with this interpretation, the levels of sustained DA over the entire cue-light period (including shock-relief and drug-reward signals) were significantly negatively correlated with punished cocaine infusions, although no correlation was observed for punished fentanyl intake. Critically, in punishment-resistant fentanyl-taking mice, we observed a pronounced increase in DA during shock delivery, indicative of a positive TD-RPE. In other words, compulsive fentanyl-taking mice may transform the aversive footshock into a highly salient sensory cue that strongly predicted drug reward. Altogether, these findings suggest that chronic drug use may disrupt normal negative TD-RPE signaling of DA neurons in NAc, thereby promoting compulsive drug taking despite negative consequences.

Although our findings provide strong support for TD-RPE as a unifying framework for interpreting NAc DA signaling underlying individual differences in excessive and compulsive drug taking, many important questions remain. We do not yet know how state value is computed, or which neural mechanisms translate these valuations into drug-seeking behaviors (e.g., probability of lever pressing), or how the observed DA dynamics in turn further modulate neural plasticity and maladaptive behaviors. Moreover, it remains unclear how different classes of drugs differentially modulate punishment sensitivity, negative RPE signaling, and the “shock-relief” rebound of DA signals. Addressing these open questions will require targeted future experiments.

### Limitations

Several limitations of the present study should be acknowledged. First, fiber photometry measures relative changes in DA release rather than absolute DA concentrations, and therefore cannot directly quantify baseline dopaminergic tone. Second, the sample size limits our ability to robustly assess sex differences in drug taking and compulsive behavior, an important factor that warrants dedicated investigation in future studies. Third, DA signaling within the nucleus accumbens is highly heterogeneous across subregions(*57*). While the present work focused on the dorsomedial shell, future studies employing more spatially resolved approaches will be necessary to systematically examine DA dynamics across distinct accumbens subregions and their contributions to addiction-related behaviors. While our computational model incorporates multiple states and agent uncertainty, we did not consider different circuit-level plasticity and maladaptive changes induced by chronic self-administration of different drugs. Furthermore, the drug-specific effect in our model is highly simplified and does not account for the myriad physiological and psychological differences in the effects of cocaine and fentanyl.

## Acknowledgments

We thank members of the Wang Lab for insightful discussions of this study. We are grateful to Priyadarshini Dutta assistance with mouse colony maintenance. This work was supported by Boston Children’s Hospital Viral Core, which is supported by NIH5P30EY012196.

## Funding

Addiction Initiative at McGovern Institute for Brain Research (FW)

The Paul E. and Lilah Newton Brain Science Award (FW)

K. Lisa Yang Integrative Computational Neuroscience (ICoN) Center fellowship (HZ)

## Author contributions

FW and KC conceived the study and designed the experiments. KC, GS, WX, CW and AS performed all experiments. KC analyzed the data. HZ and IF conceptualized and developed the computational model. KC, HZ, FW wrote the manuscript with input from IF.

## Competing interests

Authors declare that they have no competing interests.

## Data, code, and materials availability

All data and code used in the analysis are available from the corresponding authors upon request.

## Supplementary Materials

Materials and Methods

Figs. S1 to S8

## Materials and Methods

### Experimental Subjects

Adult male and female mice (C57BL/6J, 12-20 weeks old, The Jackson Laboratory) were used for this study. They were group-housed and maintained on a reversed 12-hour light/dark schedule with ad libitum access to food and water. All experimental protocols were approved by the Institutional Animal Care and Use Committee at the Massachusetts Institute of Technology.

### Surgical Procedures for Jugular Vein Catheterization

Indwelling catheters were implanted into the right jugular vein of both male and female mice, as described in the literature^53,54^. Specifically, mice were anesthetized with 1-1.5% isoflurane in oxygen (0.7 L/min) using an anesthesia mask (Part# SOMNO-0801, Kent Scientific). Once fully anesthetized, mice were placed on a heating pad (Part# 53800M, Stoelting Co.) to maintain body temperature. After shaving the hair and sanitizing the surgical area with 70% ethanol and 2% Chloroxylenol, a 2-cm mid-scapular incision was made on the back, and a second 2 cm diagonal incision was made from the right clavicle upwards to the animal’s jaw. The right jugular vein was then carefully exposed, and lifted using an Eppendorf pipette tip (Part# 13-683-718, Fisher Scientific). An 18G needle was used to create an opening in the jugular vein and a catheter (Part # C20PU-MJV1301, Instech Laboratories) was gently inserted and secured with two knots. The other end of the catheter was threaded under the skin of the shoulder to connect to a vascular access button (Part# VABM1B/25, Instech Laboratories) on the back and the incisions were sutured close. Following surgery, Mice were single-housed and received subcutaneous injections of meloxicam (5 mg/kg) daily for 2-3 days to alleviate pain and inflammation. The catheters were flushed 1-2 times daily with approximately 0.05 mL of heparinized saline (30 U/mL heparin) to maintain patency.

### Surgical Procedures for Viral Injections and Fiber Optic Cannulae Implantation

After five to seven 6-hr training sessions of cocaine or fentanyl intravenous self-administration, mice were anesthetized with 1-1.5% isoflurane in oxygen (0.7 L/min) and placed on a stereotaxic apparatus (Model 940, Kopf). A heating pad (Part# 53800M, Stoelting Co.) was used to maintain the animal’s body temperature. For viral injections, a small craniotomy was drilled above the right NAc medial shell (AP: +1.5 mm, ML:0.55 mm relative to bregma). A pulled glass pipette (Part# Q100-50-10, Sutter Instrument) front-loaded with AAV constructs (Boston Children’s Hospital Viral Core, AAV2/5-hSyn-GRAB_DA2m, 1.19E10^13^ gc/mL) was lowered into the medial shell of the NAc (DV: -4.3 mm relative to bregma). A total of 300 nL of the virus was injected at 1 nL/s with a microsyringe pump (Part# UMP3, World Precision Instruments). After the injection, the pipette was left in place for 10 minutes before being slowly withdrawn. Next, a fiber optic cannula (core diameter: 200 µm; NA: 0.37; Length: 4.5 mm, RWD Life Science) was slowly lowered to the dorsal medial shell of the NAc (∼3.7 mm below brain surface). The cannula was secured to the skull using Loctite super glue and Metabond (C&B Metabond, Parkell). The mice were allowed to recover for 4-7 days before resuming self-administration training.

### Cocaine and Fentanyl Intravenous Self-Administration (IVSA) Paradigm

One to two weeks after jugular vein catheterization surgery, the patency of implanted catheters was tested by intravenously injecting approximately 0.04 mL of a 15 mg/mL ketamine solution. Mice that passed the patency test (i.e., cessation of movement within 4 seconds) were trained to self-administer cocaine (Part# C5776, Sigma-Aldrich, 0.3 mg/kg/infusion) or fentanyl (Item# 07- 890-5657, Patterson Veterinary, 2 µg/kg/infusion) in an operant chamber (Part # ENV-307A-CT, Med Associates). The chamber was equipped with two retractable levers (Part # ENV-312-3M, Med Associates), LED lights (Part # ENV-321DM, Med Associates), and a syringe pump (Part # PHM-100VS-2, Med Associates).

Each trial of the cocaine or fentanyl IVSA started with the insertion of both levers and the illumination of the light above the active lever. Pressing the active lever triggered a drug infusion according to a fixed-ratio schedule (FR1, FR2, FR4), while pressing the inactive lever had no programmed consequences. Each infusion was followed by a 40-second time-out period during which no additional drug was delivered. This time-out period was implemented to prevent adverse health consequences associated with excessive drug intake. During the first 19.5 seconds of this time-out period, the light above the active lever blinked at 0.67 Hz (1 second on and 0.5 seconds off) with both levers remaining available, but lever pressing (i.e., time-out responses) had no programmed consequences. For the remaining 20.5 seconds of the time-out period, both levers were retracted and the lights were turned off.

Trainings began with a 3-hour auto-shaping session during which both levers were active, and pressing either lever triggered a drug infusion. In addition, a drug infusion was automatically delivered if no levers were pressed within 6 minutes. The auto-shaping session ended either when 30 infusions were delivered or after three hours had elapsed, whichever came first. Then, 6-hour long-access training sessions were followed. During these sessions, mice were trained to discriminate between an active drug-delivering lever and an inactive lever. Only presses on the active lever resulted in drug infusions. The active lever was designated as the non-preferred lever, based on behavior observation during the initial auto-shaping session. To prevent the catheter blockage during the 6-hour training sessions, automatic drug infusions were delivered if the active lever was not pressed within 30 minutes. To prevent adverse health effects of excess drug intake, the maximum number of infusions per session was capped at 150 for males and 120 for females. The training protocol consisted of 7-9 sessions on an FR1 schedule, followed by 2 sessions on an FR2 schedule, and 10 sessions on an FR4 schedule. Mice were trained 5 days per week.

Following this long-access training, drug-taking behavior and fiber photometry recordings were conducted during 3-hour IVSA sessions under an FR4 schedule. The animal’s behavior was also videotaped. Mice completed at least three 3-hour baseline IVSA sessions before undergoing three consecutive punishment sessions. During these punishment sessions, each drug infusion was paired with a brief, mild foot shock (Intensity: 0.2 mA; duration: 0.5 seconds). The shock intensity was verified using an ammeter (ENV-420, Med Associates) before each punishment session. After completion of the punishment sessions, catheter patency was tested prior to brain tissue collection, and only mice that pass this test were included in the final analysis of cocaine or fentanyl IVSA. In total, 24 out of 27 mice successfully completed cocaine IVSA training and testing with confirmed catheter patency, and 27 out of 29 mice completed the fentanyl IVSA training and testing with confirmed patency. As a control, 8 mice completed the saline IVSA training and testing, six of which underwent fiber photometry recordings.

### Fiber Photometry

Dopamine transmission in the medial shell of the NAc was recorded with a rotary fiber photometry system (Part# RFPS_2S_GCaMP_RedFluo, Doric). The system was equipped with an assisted electrical rotary joint (Part# AHRJ-EL_24_FMC_25, Doric) for fiber photometry and a fluid rotary joint for infusing drugs. As the optic path and light detector of the photometry system were integrated as one small device (i.e., rotary fluorescence mini cube), which rotated as mice move in the chamber, the fiber bending and movement-induced artifacts were minimized. To measure signals from the green DA sensor^49,50^ GRAB_DA2m, purple (405 nm) and blue (465 nm) LEDs within the fluorescence mini cube (RMFM, Doric Lenses) emit a sinusoid illumination at 208.616 Hz and 572.205 Hz respectively to excite the fluorophore. The power at the tip of the patch cable was 5-10 µW. A 0.4-meter low auto-fluorescence fiber optic patch cord was used to connect the mini cube to the implanted fiber optic cannulae. Bulk fluorescent signals were detected with detectors integrated within the mini cube, amplified by a Doric fluorescence detector amplifier, and digitized at 12k Hz by a fiber photometry console (FPC, Doric Lenses) which also recorded behavioral events of drug infusion, cue presentation and lever presses from Med Associates operant chamber. The digitized signals were lock-in demodulated based on the frequency of excitation lights (405nm and 465 nm) and down-sampled to 120 Hz. Doric Neuroscience Studio was used to acquire and stream demodulated signals to the disk. To diminish photobleaching, the fiber photometry system was automatically turned ON for 30 minutes and then turned OFF for 30 minutes. This ON-and-OFF cycle automatically repeated 3 times to cover the entire 3-hour IVSA testing phase.

### Histological Staining

Mice were deeply anesthetized with isoflurane and intracardially perfused with 1x PBS followed by 4% paraformaldehyde. The brain was post-fixed overnight with 4% paraformaldehyde and cryo-protected with 30% sucrose for 2-3 days. The brain was then cut with a cryostat into 80 μm coronal slices. For visualizing canula tracks and the expression of GRAB_DA2m, slices were stained with DAPI (1:5000 dilution, H3570, ThermoFisher, Waltham, MA) or fluorescent Nissl stain (1:500 dilution, N21479, ThermoFisher).

### Data Analysis

Data analysis was performed using Doric Neuroscience Studio and custom scripts written in Python and MATLAB (MathWorks, Natick, MA).

### Behavior analysis during the IVSA

The total number of drug infusions, active lever presses and inactive lever presses were recorded. In addition, the timestamps of trial onset and all behavioral events were recorded as well. These timestamps were used to generate raster plots of lever-press activity relative to the trial start.

To classify mice as low or high drug takers, the average number of infusions per mouse during the 3-hour baseline IVSA sessions was calculated. Mice with the average baseline infusion counts greater than the group mean plus 10% of the mean were classified as high drug takers, whereas mice with average baseline infusion counts lower than the group mean minus 10% of the mean were classified as low drug takers. Mice that fell between these thresholds were left ungrouped.

Similarly, to classify mice as high or low punishment-resistant, the average number of punished infusions per mouse during the 3-hour punishment sessions was calculated. Mice with the average punished infusion counts greater than the group mean plus 10% of the mean were classified as high punishment-resistant, whereas mice with average punished infusion counts lower than the group mean minus 10% of the mean were classified as low punishment-resistant. Mice that fell between these thresholds were left ungrouped.

### Fiber photometry data analysis

The fiber photometry data were preprocessed using Doric Neuroscience Studio to extract the Z-score of the DA dynamics based on a published method^55^. Specifically, signals recorded at both 465 nm and 405 nm (i.e., isosbestic signal) were smoothed by applying a running average with a window size of 0.1 seconds. The bleaching slope and low-frequency fluctuations of both signals were corrected with an adaptive, iterative re-weighted penalized least squares algorithm^56^. Both signals were standardized by calculating their Z-scores. Non-negative robust linear regression was used to fit the Z-score of signals at 405 nm to those at 465 nm. Finally, DA dynamics is calculated by subtracting the fitted Z-score of signals at 405 nm from the Z-score of the signals at 465 nm.

Drug self-administration-evoked DA responses (hereafter referred to as “drug-evoked” responses) were analyzed by aligning DA dynamics to the onset of each drug self-administration and constructing peri-event time histograms (PETHs). Importantly, the generation of drug-evoked events required response contingency, cue presentation, and simultaneous drug delivery. Normalized PETH was obtained by averaging PETHs across trials and subtracting baseline DA activity (-10 to 0 s relative to drug infusion). Onset DA responses were defined as the mean evoked DA signals within 1 s of drug infusions, whereas sustained DA responses were defined as the mean evoked DA responses from 1 to 19.5 s post-infusion (corresponding to drug-associated cue period).

To analyze DA responses to active lever presses, DA dynamics were aligned to the first active lever press in each trial to construct PETHs. The PETHs were then normalized by averaging PETHs across all trials and subtracting the mean baseline activity (the 2-s interval immediately preceding the press). DA responses to active lever press were quantified as the mean evoked DA signals within the 2 s following the active lever press.

To analyze the decay rate of DA transients, DA dynamics from baseline sessions were first low-pass filtered using a 4^th^-order Butterworh filter. Local peaks and their subsequent troughs were then extracted with the *findpeaks* function in MATLAB. Finally, we performed a linear regression on these corresponding peaks and troughs, and the slope of the regression was defined as the decay slope.

### Temporal Difference Learning in Actor-Critic Model

In TD-learning considering self-administration, an agent transitions through a sequence of states *S*(*t*) according to its policy *M* interacting with a Markov decision process (or a semi-Markov decision process). The ‘Critic’ computes the value associated with each state, defined as the expected discounted future returns based on current policy *M*:

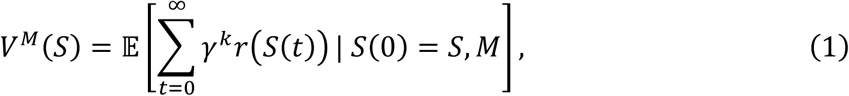

where *t* denotes time and *S*(*t*) is the state visited at time *t*. *r*(*S*(*t*)) denotes the reward delivered at state *S*(*t*), and *γ* ∈ (0, 1) is a discount factor. In the experiments we examine, the drug reward is present with delay after infusion event and lasts for a prolonged period until the end of the trial. The ‘Actor’ aims to learn an optimal policy *M*^∗^ that maximizes the expected total future returns:

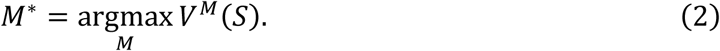

Interleaved steps of estimating the value function and updating the policy are used in learning. For estimating the value function, under the Markov property, the value at time t for state *S*(*t*) can be rewritten as a sum of the reward received at *t* and the discounted value at the next time step:

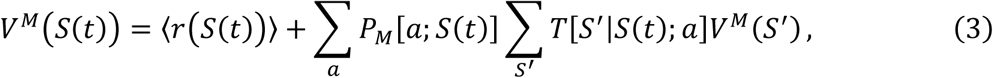

where *P*_*M*_[*a*; *S*(*t*)] denotes the probability of choosing action *a* at state *S*(*t*) according to the policy *M* and *T*[*S*^′^|*S*(*t*); *a*] denotes the probability of state transition from *S*(*t*) to *S*(*t* + 1) = *S*^′^ at next time step *t* + 1 when taking action *a*. 〈*r*(*S*(*t*))〉 denotes the mean reward received at *S*(*t*). Temporal difference learning takes *r*(*S*(*t*)) + *V*^*M*^(*S*′) as a Monte Carlo sample to approximate the right side of equation (3) and then bootstrap by replacing the unknown *V*^*M*^(*S*′) with the current estimate *V*(*S*′). Therefore,

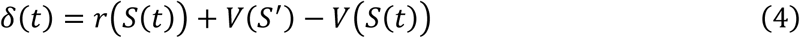

is used as a sampled approximation to the mismatch *V*^*M*^(*S*(*t*)) − *V*(*S*(*t*)). *δ*(*t*) is called temporal-difference reward prediction error (TD-RPE). When *δ*(*t*) = 0, the value function is well approximated. However, when *δ*(*t*) is positive or negative, the Critic’s estimate *V*(*S*(*t*)) should be increased or decreased, respectively (*ɑ* is learning rate):

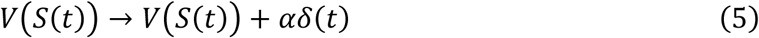

For updating the policy to satisfy equation (2), we could similarly use *r*(*S*(*t*)) + *V*(*S*^′^) as a Monte Carlo estimates of right side of equation (3), and the policy is updated along the gradient of equation (X3). If *P*_*M*_[*a*; *S*] is parameterized as a SoftMax distribution:

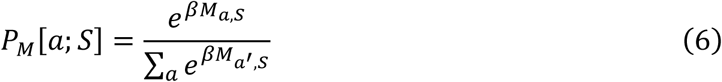

where *M*_*a*,*S*_ denote the action value for taking action *a* at state *S*, and *β* is the SoftMax parameter, then the update of Actor’s policy upon taking action *a* at state *S* has an elegant and bio-plausible formular:

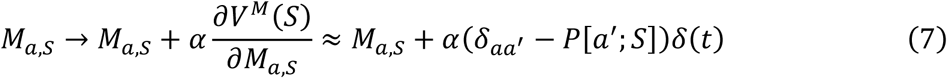

where *δaa*′ = 1 if *a*^′^ = *a* and 0 otherwise. *δ*(*t*) is the RPE defined above and *ɑ* is the learning rate.

### Internal state inferred from task stimulus

For simplicity, we assume each trial is decomposed into 3 discrete internal state according to different sensory feedback:

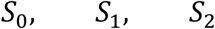

*S*_0_denotes the initial trial stage from light-onset and lever-insert to drug-infuse, which dominates the trial period. *S*_1_denotes the 20s delay period from drug-infuse to lever-retract, with blinking light on and off periodically. *S*_2_ denotes the 20s light-off ending-stage of each trial, from lever-retract to next lever-insert. While the external task progression is governed deterministically by the external clock and lever-press counter, the animal does not have access to either external clock or counter and stochastically transit among these states based on sensory input generated at each external task stage.

### Internal state triggered by shock

Foot shock is a life-threatening stimulus to animals and usually triggers leaping, retreating, scanning, or freezing behaviors. After a period of alert, the animal gradually returns to the spontaneous behavior as pre-shock stage, suggesting a process of inferring what is happening. Therefore, we assume the shock triggers another cascade of internal states:

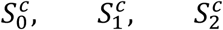

where 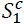 denotes the alert period once getting the shock. After scanning the surroundings, the shock state transit to 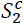 stochastically, where 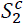 denotes the ‘safety’ state. 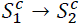 corresponds to shock relief. Once the light is turned on again for next trial, the safety state transit to 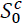, which denotes the ‘danger’ state, extending from lever-insert to getting-shock. A negative reward *r*^*c*^ < 0 is provided during transition 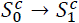 and an innate negative value 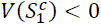 is initially assigned to shock state 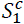. Experimental data for baseline shock testing confirms that first shock can evoke lasting DA dynamics independent of drug reward, rendering additionally considering shock-related states necessary.

Given both task-related states *S*(*t*) and shock-related states *S*^*c*^(*t*), the total value at time *t* is a linear combination of the state value and action value:

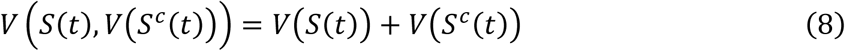

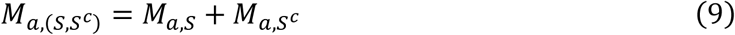

All the variables *V*(*S*(*t*)), *V*(*S*^*c*^(*t*)), *Ma*,*S*, *Ma*,*Sc* follow the TD-learning rule for Actor-Critic model introduced above.

### Uncertainty in state transition during long-access training

Each task trial has an extended duration, lasting 2-3 mins on average, which far exceeds the experimental timescale for classical conditioning in reinforcement learning. This may cause several non-negligible outcomes on animal’s behavior. First, due to constant spontaneous movement during operant stage, the animal may not detect a transient cue and cue-evoked internal state transition can be delayed. For example, the transition *S*_0_ → *S*_1_ may lag behind the first light-blink, which is consistent with the trial-to-trial jittered initial DA response timing. For this reason, we assume that the animal has a probability to transition from *S*_0_to *S*_1_ at each light blink (from light on to light off):

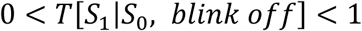

Second, during the ON cycle of blinking light after infusion, the animal has a non-zero probability to return to ground state *S*_0_. This is because *S*_0_ occupies the majority of trial period with light-on as sensory input, therefore the animal may treat *S*_0_ as the “ground state” with large prior. Considering that the ON cycle of blinking light during *S*_1_ shares the same sensory input as *S*_0_, the animal may switch its internal state to *S*_0_ during on-edge and return back to *S*_1_ at next off-edge of blink, which we call “diffuse”

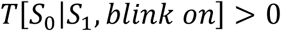

### Individual difference on action cost and shock sensitivity

In the model, we only consider binary action for simplicity: *a* = 0 denotes no-press while *a* = 1 denotes lever-press. We assume that the cost for lever-press varies across animals. Therefore, when updating the action value for pressing lever, an additional action cost *c*(*a*) is considered:

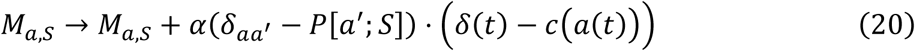

where *δ*(*t*) − *c*(*a*(*t*)) = (*r*(*t*) − *c*(*a*(*t*))) + *V*(*S*(*t* + 1)) − *V*(*S*(*t*)) is the combined RPE for Actor. Variations in action cost has profound impact on animal’s addiction behavior: for example, we find that animals who learned to press lever by biting onto the lever, or using the chin to press lever, had significantly more lever press counts than those pressing by using the paw, and consequently more drug taking. Indeed, pressing the lever with a paw is relatively demanding: the animal must rear up, maintain balance, and then lift a paw to make a press. During shock session, we find that some animals show much higher tolerance to the electric shock compared to others. Therefore, shock sensitivity is another factor underlying individual difference.

### Effects of Cocaine and Fentanyl on modulating shock sensitivity

Drugs like cocaine and fentanyl have diverse pharmacological effects on animals besides serving as reinforcers. Specifically, cocaine is a psychostimulant, whereas fentanyl is an opioid analgesic. Therefore, we assume that the two drugs modulate shock sensitivity in opposite directions: cocaine lowers the shock threshold, whereas fentanyl raises it.

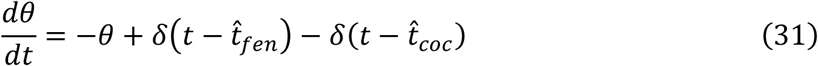

where 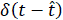 is a Dirac delta that represents an event occurring at time 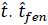 and 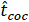 denote the infusion event of fentanyl and cocaine respectively. *θ* is the shock threshold for computing effective shock reward *r*^*c*^:

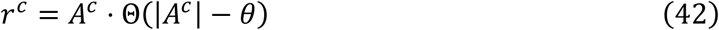

where *A*^*c*^ is a measure of shock strength, and Θ is step function.

### Cocaine’s effect on DA signal decay

Since cocaine blocks dopamine reuptake, extracellular DA clears more slowly, extending the decay timescale relative to normal condition; in our recordings, DA decays roughly fivefold more slowly than normal. In the cocaine model, we represent each RPE event as producing a DA transient with a slow decay. Importantly, the DA level during this decay is not treated as additional teaching signal: only the phasic RPE at the event time drives learning. We instead interpret the slowly decaying component as a motivational modulation signal, consistent with its timescale.

---

**Fig. S1.**
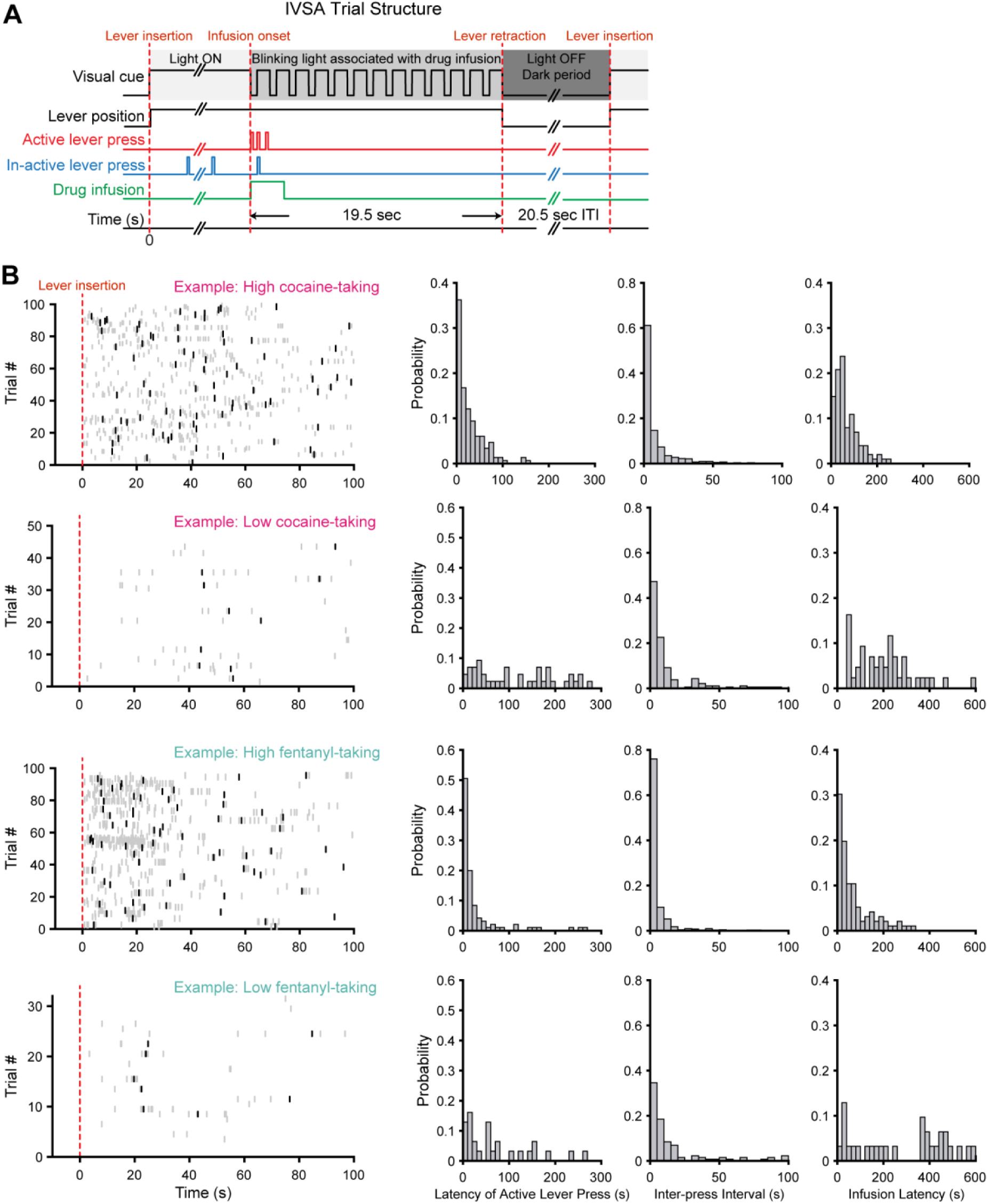
Drug-taking behavior. (**A**) Schematic of the drug self-administration task structure (showed with FR1 schedule), highlighting the light-ON, light-blinking, and light-OFF epochs. (**B**) Raster plots of active lever presses (gray lines) and infusions (black lines) aligned to the start of each trial (lever insertion; time 0), along with distribution of the latency to the first active lever press, inter-press interval and infusion latency relative to trial onset. Each row corresponds to an example mouse.

**Fig. S2.**
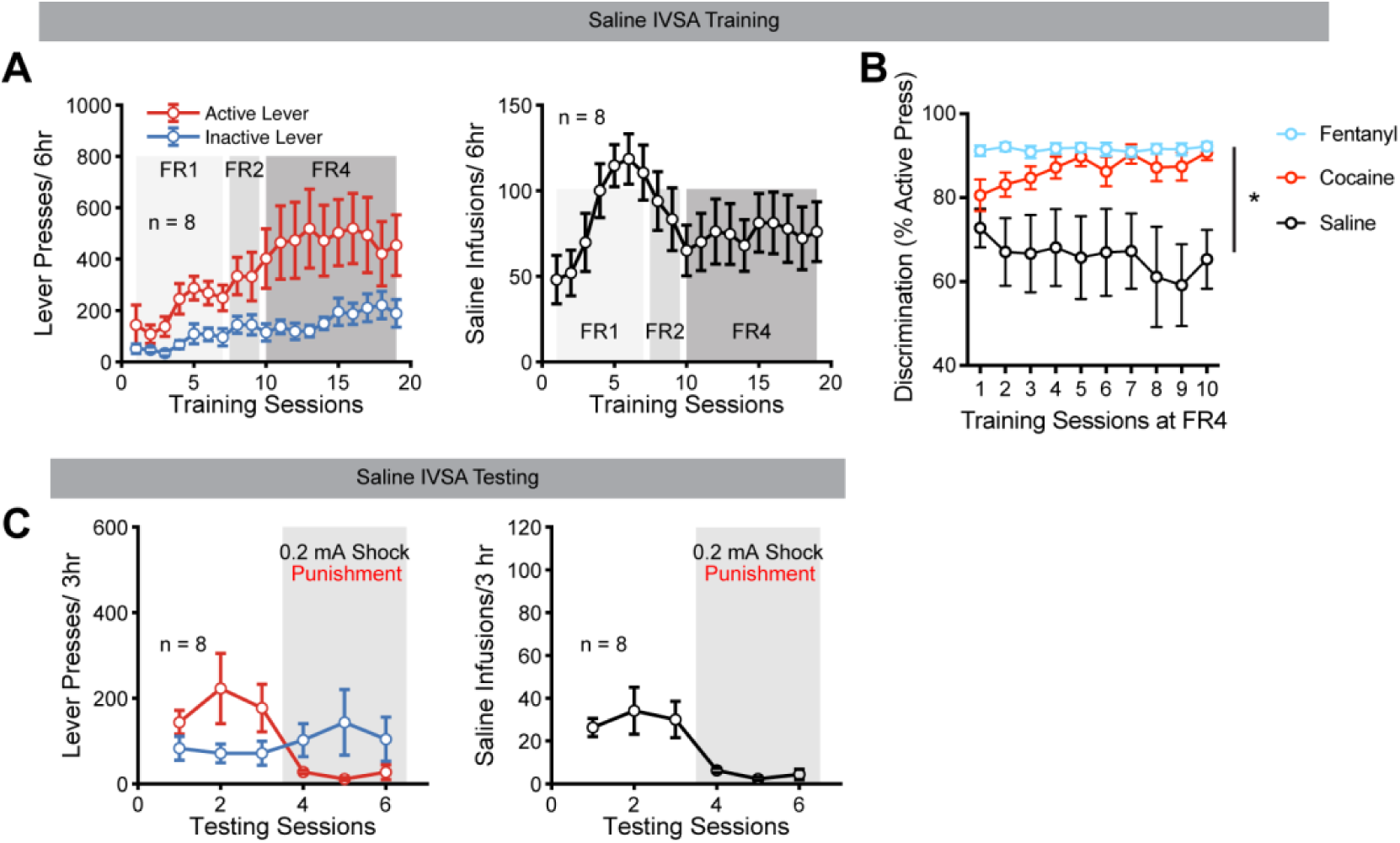
Saline intravenous self-administration (IVSA) behaviors. (**A**) Lever presses (left) and saline infusions (right) across daily 6-hour saline IVSA training sessions (n = 8). (**B**) Lever discrimination (proportion of active lever presses) across 10 training sessions of cocaine (red), fentanyl (blue) and saline (black) IVSA under FR 4 schedule (repeated two-way ANOVA test). (**C**) Lever presses (left) and saline infusions (right) during the testing phases of saline IVSA (n = 8). Error bars indicate mean ± standard error mean (SEM). * represents p < 0.05.

**Fig. S3.**
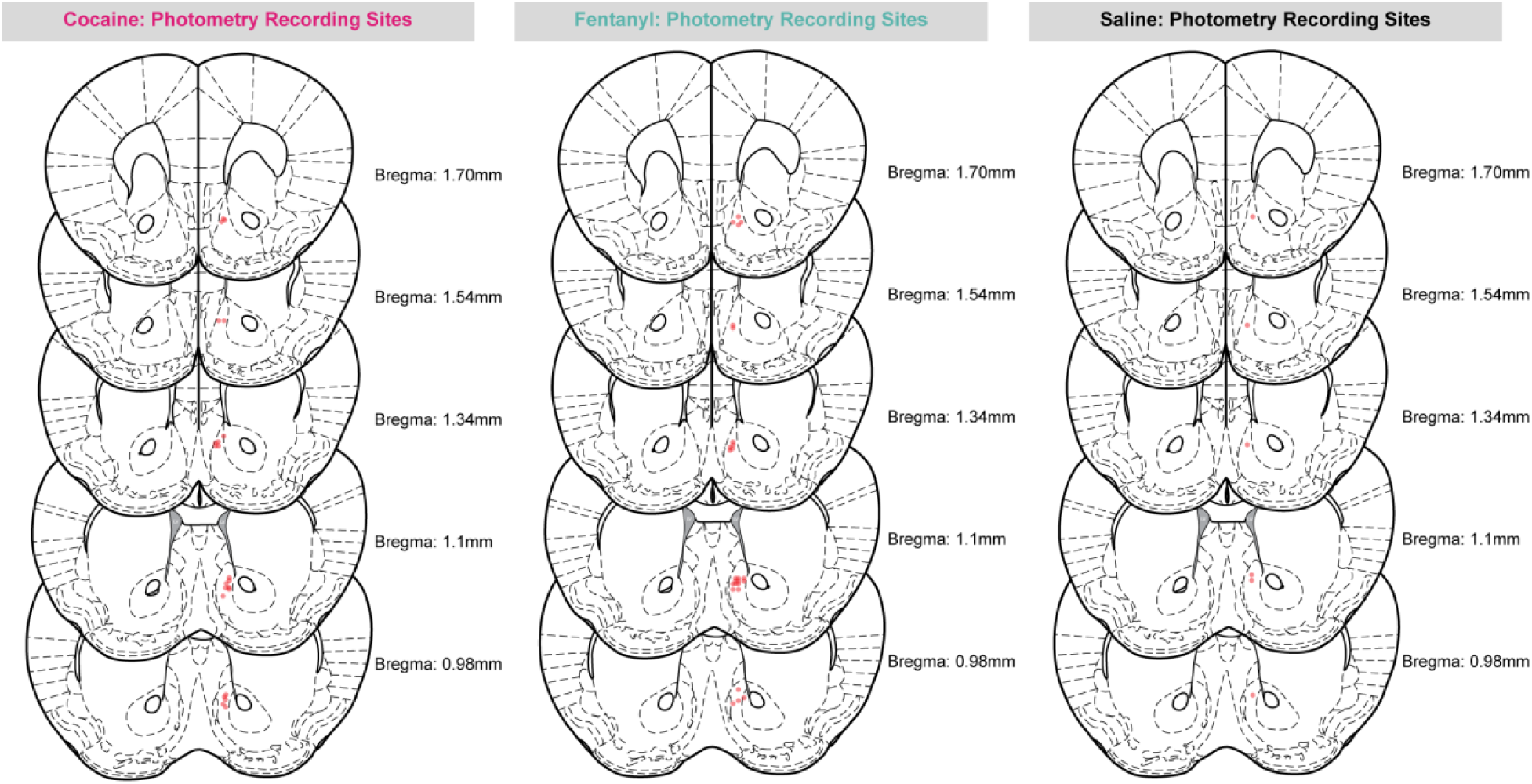
Histological verification of optical fiber placements. The tips of optical fiber tracks are indicated by pink circles (one circle per mouse). Coronal sections are labeled with anterior-posterior coordinates relative Bregma. In some cases (n= 2), placements could not be verified because the fiber tracks were not detectable.

**Fig. S4.**
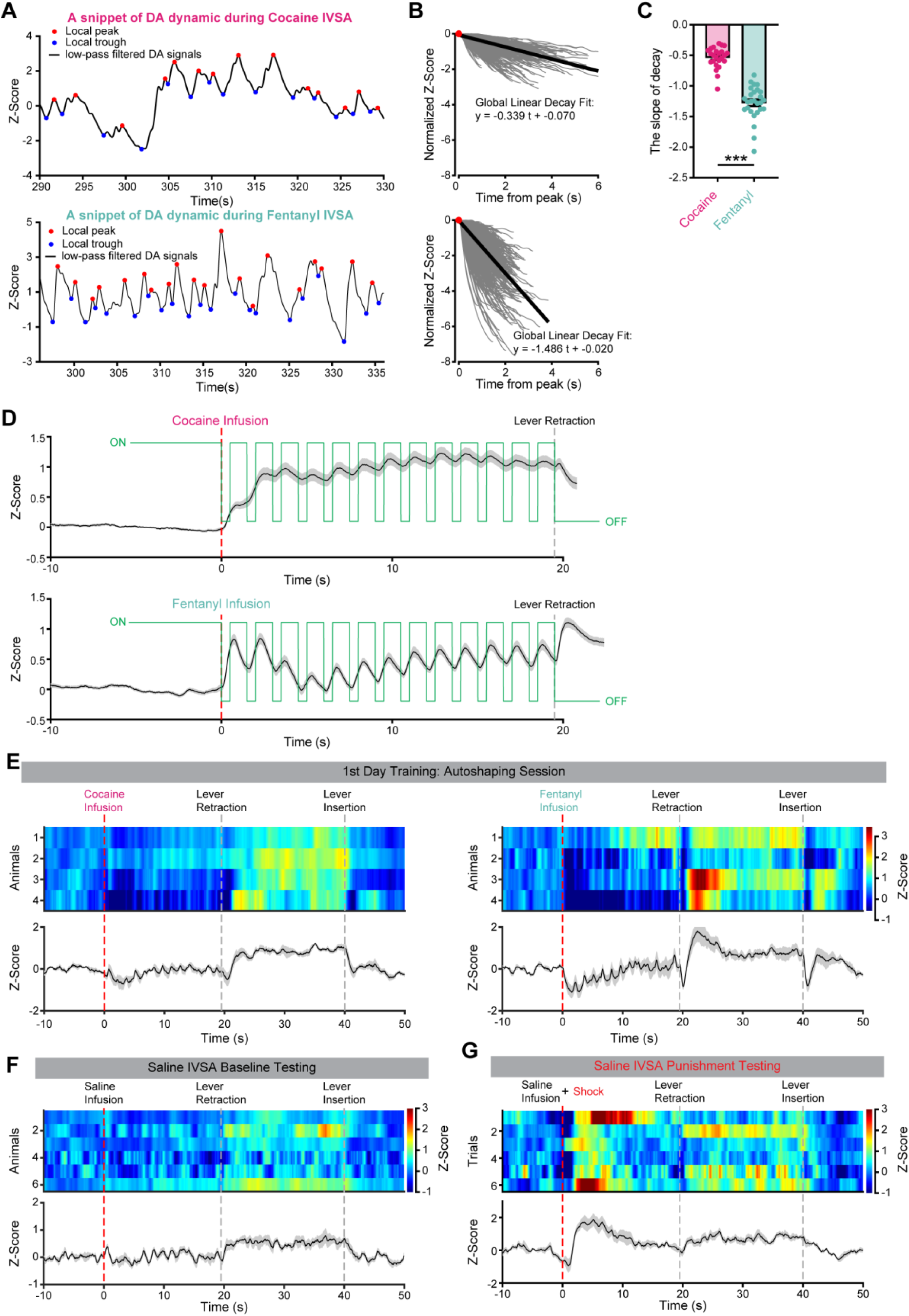
Dopamine dynamics during cocaine, fentanyl, and saline IVSA. (**A**) Example traces of low-pass filtered, z-scored dopamine signals. Local peaks (red) and troughs (blue) are indicated. (**B**) Linear regressions fitted to decay segments from each local peak to the subsequent trough, corresponding to the example traces showed in (**A**). (**C**) Bar plots of decay slopes in mice IVSA cocaine and fentanyl (Welch’s t-test). (**D**) Population-averaged dopamine responses to cocaine (top) and fentanyl (bottom) IVSA overlaid with the blinking light signal (green line). (**E**) Dopamine responses to cocaine or fentanyl IVSA on the 1^st^ day of training. (**F**) Dopamine responses to saline IVSA during the 3-hour testing phase after extended training. (**G**) Dopamine responses to the co-occurrence of saline IVSA and footshock during punishment sessions of saline IVSA. Error bars indicate mean ± standard error mean (SEM). *** represents p < 0.001.

**Fig. S5.**
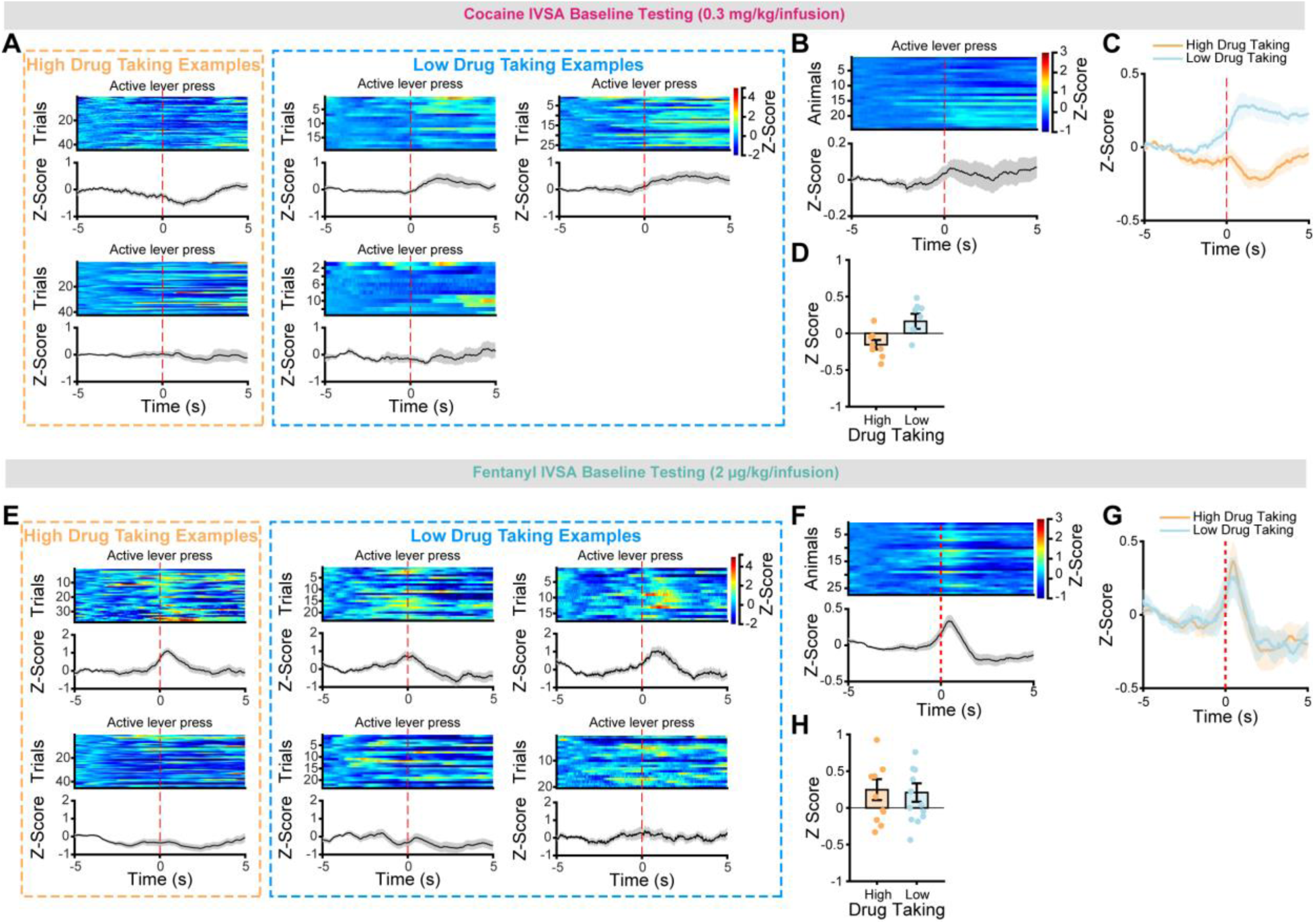
Dopamine responses to active lever presses. (**A**) Examples dopamine responses to the first active lever press of each trial. Two examples are from high cocaine-taking mice and three from low cocaine-taking mice. (**B**) Group dopamine responses to active lever presses. Each row of the colormap represents one mouse, sorted by the number of infusions (top = most). (**C**) Time courses of DA responses to active lever press for high (orange) and low (light blue) cocaine-taking mice. (**D**) Modulated dopamine responses by active lever presses for high (orange) and low (light blue) cocaine-taking mice. Responses were calculated as the difference between the mean z-scored dopamine signals from 0-2 s after the active lever press and the -2 to 0 s baseline before the press. (**E-H**) Same analyses as (**A-D**), but for fentanyl IVSA.

**Fig. S6.**
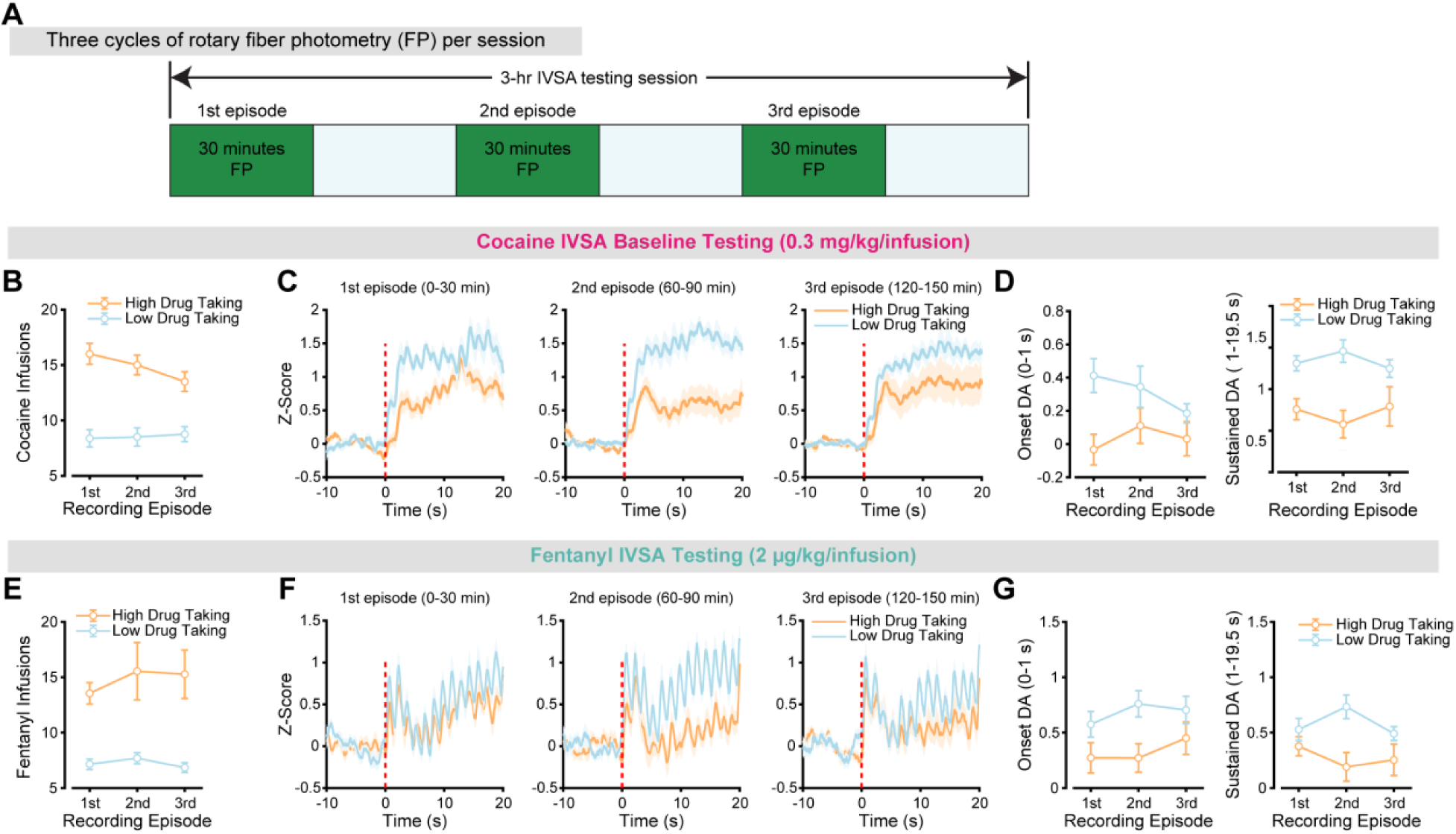
Dopamine responses across three episodes of recordings during cocaine and fentanyl IVSA. (**A**) Schematic illustrating three cycles of rotary fiber photometry during 3-hour IVSA testing sessions. (**B**) Number of cocaine infusions across the three recording episodes during 3-hour IVSA. (**C**) Time courses of DA responses to contingent cocaine infusions for high (orange) and low (light blue) cocaine-taking mice across the three recording episodes. (**D**) Averaged onset and sustained DA responses across the three recording episodes for high (orange) and low (light blue) cocaine-taking mice. (**E-G**) Same analyses as in (**B-D**), but for fentanyl.

**Fig. S7.**
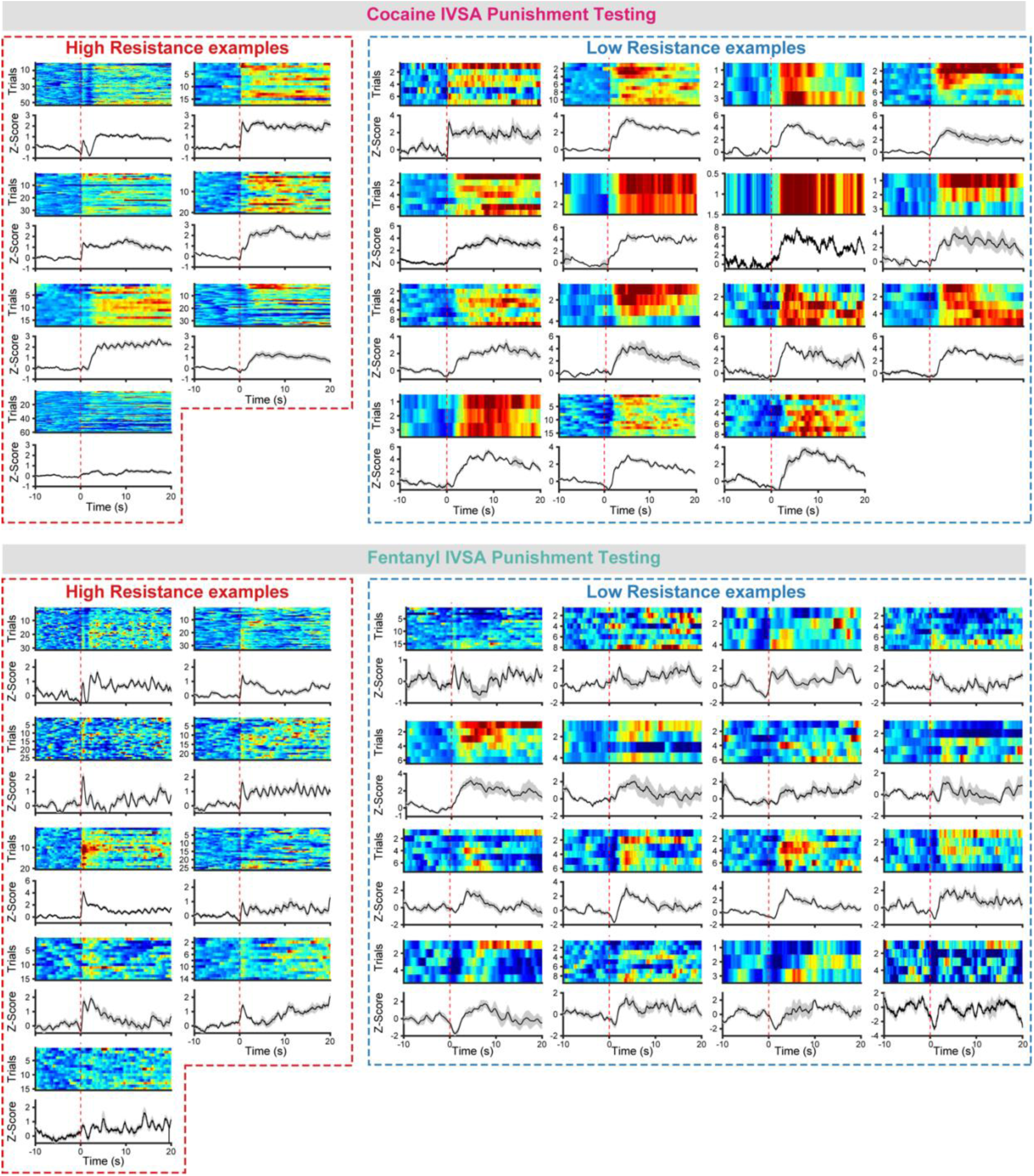
Dopamine dynamics during punished drug taking in individual mice. Each colormap represents a single mouse. Dash lines at time 0 represent the onset of drug infusion plus a mild foot shock (0.5 s).

**Fig. S8.**
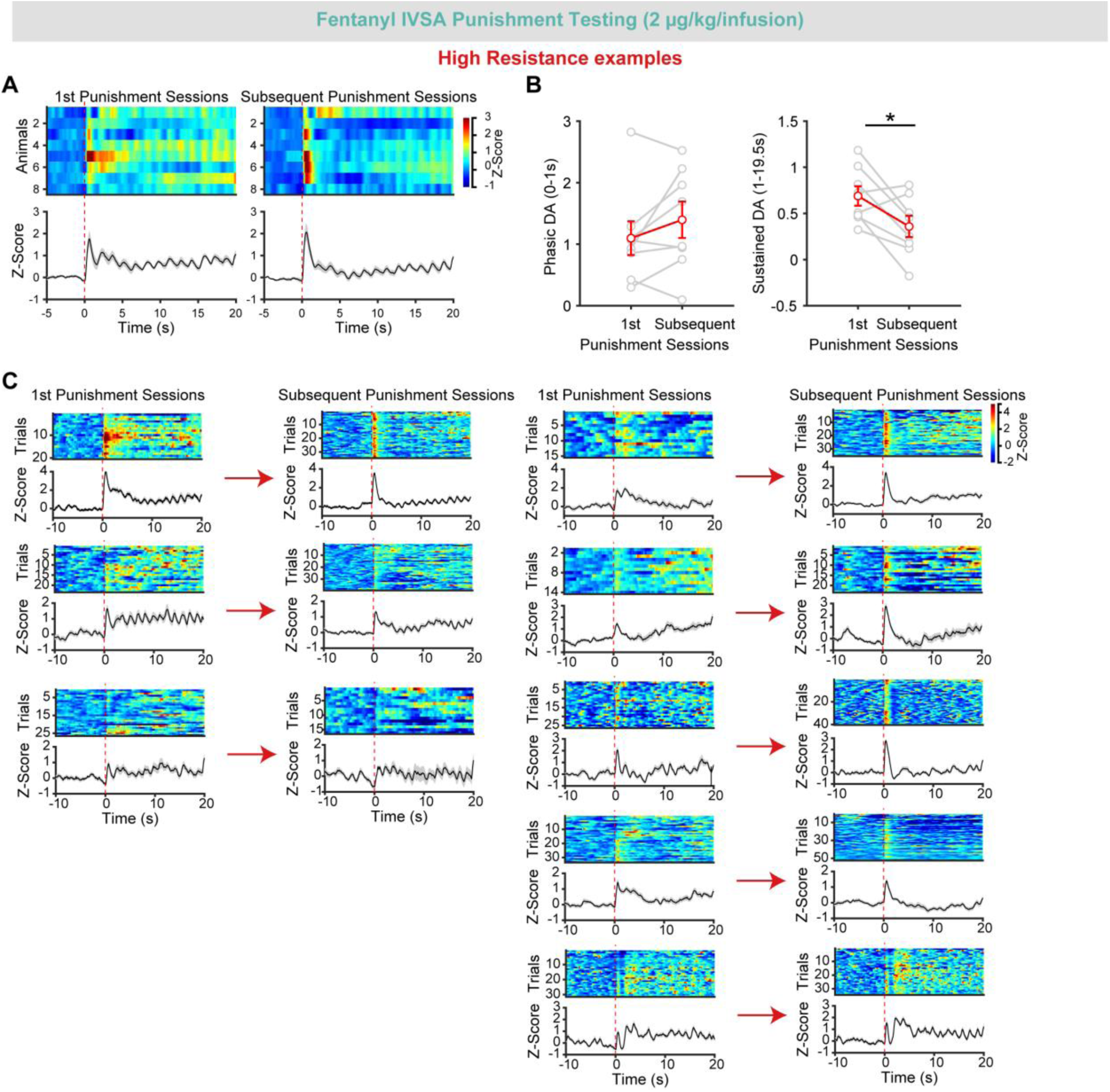
Changes in dopamine dynamics across punishment sessions of fentanyl IVSA. (**A**) Group DA responses to punished fentanyl infusions in punishment-resistant mice (n = 8) during the 1^st^ punishment session (left) and a subsequent punishment session (right). Each row of the colormap represents the same mouse recorded across punishment sessions. (**B**) Quantification of onset and sustained DA responses during the 1^st^ and subsequent punishment sessions. (**C**) Individual examples of DA responses across each trial during the 1^st^ and subsequent punishment sessions. * represents p < 0.05.

## Notes

### Competing Interest Statement

The authors have declared no competing interest.

